# Opposing neural signals of Ca_V_1-encoded peptides are tuned by C-terminus mediated inhibition

**DOI:** 10.1101/2020.01.26.920348

**Authors:** Yaxiong Yang, Min Liu, Nan Liu, Wenxiang Li, Zhen Yu, Weili Hong, Ping Li, He Jiang, Haiyan Ge, Ping Wang, Sen Song, Xiaomei Li, Yubo Fan, Xiaodong Liu

## Abstract

L-type calcium (Ca_V_1) channels regulate gene expressions via the cascade of excitation-transcription coupling, or directly as standalone CCAT (Calcium Channel Associated Transcriptional-regulator) peptides encoding distal carboxyl-terminus (DCT) of Ca_V_1, both evidenced in dendritogenesis signaling in neurons. We here discover that DCT peptides opposedly mediate these two sets of transcription signals, all tunable in accordance to C-terminus mediated inhibition (CMI) of Ca^2+^/Ca_V_1 influx. By electrophysiology, neurite morphology, and FRET 2-hybrid binding analyses, we systematically examined native and derived DCT peptides across Ca_V_1, unveiling that the overall balance between cytosolic inhibition *versus* nuclear facilitation is spatially and temporally tuned by CMI of each DCT variant. Our findings not only resolve several controversies existing to DCT variants, but also propose a *de novo* scheme of Ca_V_1-centric gene regulation: two concurrent routes of transcription signals initiated from either membrane Ca_V_1 channels or nuclear Ca_V_1-encoded peptides are subject to autonomous feedback tuning by peptide/channel interactions.

## Introduction

Voltage-gated Ca^2+^ channels (Ca_V_) generate Ca^2+^ signals triggering diverse events of pathophysiological significance, in response to electrical potentials across the cell membrane (Catterall 2011). L-type Ca^2+^ channels (Ca_V_1) of Ca_V_ family, especially Ca_V_1.2 and Ca_V_1.3, are widely expressed in human tissues and organs including the nervous system (Simms and Zamponi 2014). Among its multiple roles manifested in various biological processes, Ca_V_1 channels mediate the excitation-transcription coupling which transduces transmembrane Ca^2+^ into nuclear signals to regulate transcription and expression of multiple genes, including those important to morphological and functional plasticity. Such cytonuclear signaling cascade is coordinated by multiple molecules including calmodulin (CaM), Ca^2+^/CaM-dependent Kinase II (CaMKII) and calcineurin, altogether to tune channel gating and Ca^2+^ influx, cytonuclear translocation of key molecules/signals, activate nuclear transcription factors such as CREB and NFAT, and regulate gene expression for neuronal function and development (Murphy, Sanderson et al. 2014, Li, Tadross et al. 2016). Intriguingly, in addition to cytonuclear excitation-transcription coupling mediated by the full-length channel, the peptide fragments encoded by distal carboxyl-terminus (DCT) of Ca_V_1 could also act as transcription factors (TFs) in the nucleus that directly regulate gene expression to promote neurite outgrowth (Redmond, Kashani et al. 2002, Wong and Ghosh 2002, Gomez-Ospina, Tsuruta et al. 2006). Meanwhile, DCT has been reported to acutely inhibit Ca^2+^ entry into the cell (C-terminus mediated inhibition or CMI), by competing against apoCaM (Ca^2+^-free form of CaM) and thus concurrently inhibiting both voltage-gated activation (VGA) and Ca^2+^-dependent inhibition (CDI) (Liu, Yang et al. 2017). It is postulated that CMI should downregulate Ca^2+^/Ca_V_1 signaling along the cascade of excitation-transcription coupling in neurons, in that the conventional inhibitor dihydropyridine (DHP) indeed tunes down Ca_V_1-dependent transcriptional signaling (Redmond, Kashani et al. 2002, Wheeler, Groth et al. 2012). However, if the above hypothesis gets proved, the actual roles of DCT would encounter an apparent dilemma: promotion of neurite outgrowth by regulating related genes as nuclear TF, in direct opposition to inhibition of Ca^2+^ influx and presumably also neurite outgrowth as channel inhibitor. In this work, we undertook the mission to elucidate the above controversy regarding Ca_V_1-encoded peptides.

In neurons, there are two independent sources of peptides encoding DCT fragments. One is from the activation of a cryptic promoter in the coding region of CACNA1C (Ca_V_1.2). Such DCT fragment is termed as calcium channel associated transcriptional regulator (CCAT_C_) and contains intact proximal C-terminal regulatory domain, nuclear retention domain and distal C-terminal regulatory domain (PCRD, NRD and DCRD, respectively) (Figure 2-figure supplement 1) (Gomez-Ospina, Panagiotakos et al. 2013). Here NRD is an indispensable region for Ca^2+^-dependent nuclear transport (Gomez-Ospina, Tsuruta et al. 2006), whereas PCRD and DCRD serve as key elements for CMI (Liu, Yang et al. 2017). Two putative transcription activation domains of DCT fragments are distributed on PCRD-NRD junction and DCRD (Gomez-Ospina, Tsuruta et al. 2006), the latter of which appears to have more contributions (Gomez-Ospina, Panagiotakos et al. 2013). Alternatively, DCT peptides could be generated from proteolytic cleavage of the full-length Ca_V_1. The cleavage sites of amino acid sequences’NNAN’on DCT are highly conserved from Ca_V_1.1 to Ca_V_1.3 (Hulme, Konoki et al. 2005). Cleaved C-termini of Ca_V_1.2 (CCT_C_) and Ca_V_1.3 (CCT_D_) have only incomplete PCRD domains, while still able to function as TFs to regulate gene expressions due to their intact NRD and DCRD (Schroder, Byse et al. 2009, Lu, Sirish et al. 2015).

**Figure 1.**
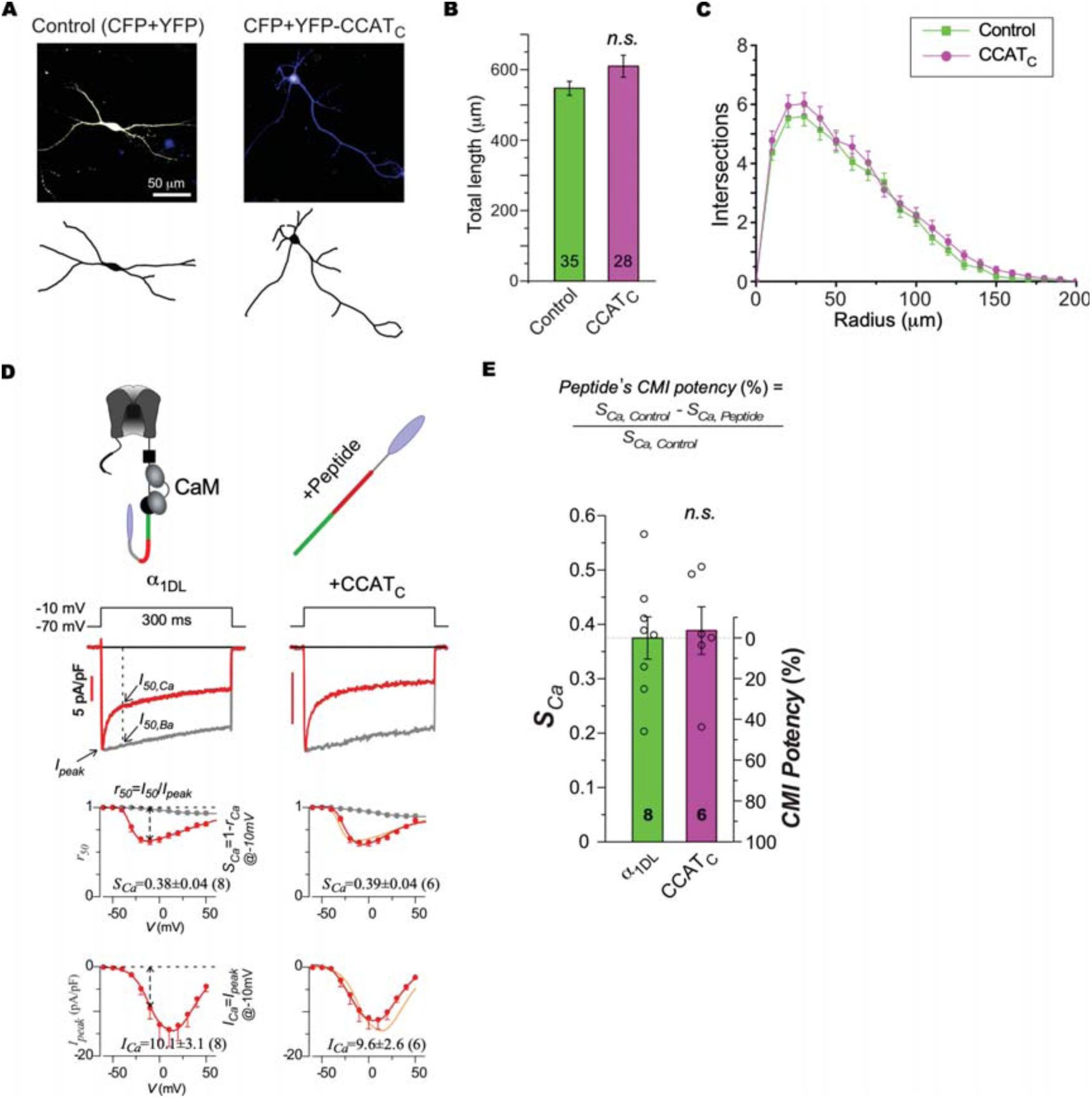
Ca_V_1.2-encoded CCAT_C_ exhibited neither TF nor CMI capabilities in cortical neurons. (A) Representative confocal fluorescence images via merged CFP/YFP channels (upper) and neurite tracings (lower) for cultured cortical neurons (DIV-7), transfected with YFP or YFP-CCAT_C_, respectively. CFP was co-transfected for both groups to trace neurite morphology. (**B** and **C**) Statistical summary of total neurite length (**B**) and *Sholl* analyses (**C**), with the total numbers of cells indicted in parentheses. (**D**) Effects of CCAT_C_ on basic channel functions of Ca_V_1.3 (α_1DL_) heterologously overexpressed in HEK293 cells, illustrated at the top row. Representative Ca^2+^ current (trace with scale bar in red) and Ba^2+^ current (rescaled in gray) were elicited by voltage step at −10 mV (the second row to the top). The next two rows show the inactivation and activation profiles, respectively with *r_50_* (ratio between current at 50 ms and peak current) for inactivation and *I_peak_* (Ca^2+^ current) for activation across the full range of membrane potentials (*V*). *S_Ca_*, quantified by 1-*r_50,Ca_* at −10 mV, serves as the major index for inactivation (CDI), whereas *I_Ca_*, quantified by *I_peak_* at −10 mV, serves as the index of activation (VGA). The groups of CCAT_C_ (right column) and control (left column, also replicated in light red) were compared for their inactivation (*r_50_*) and activation (*I_peak_*) profiles. (**E**) Statistical summary of *S_Ca_* and peptide’s CMI potency. To evaluate the potency of the DCT peptide to attenuate inactivation (*S_Ca_*), CMI potency is defined as the radiometric parameter (in percentage): (*S_Ca,control_*-*S_Ca,exp_*)/*S_Ca,control_*. Student’s *t*-tests were used for and (**E**). Data were shown as means±SEM.

**Figure 2.**
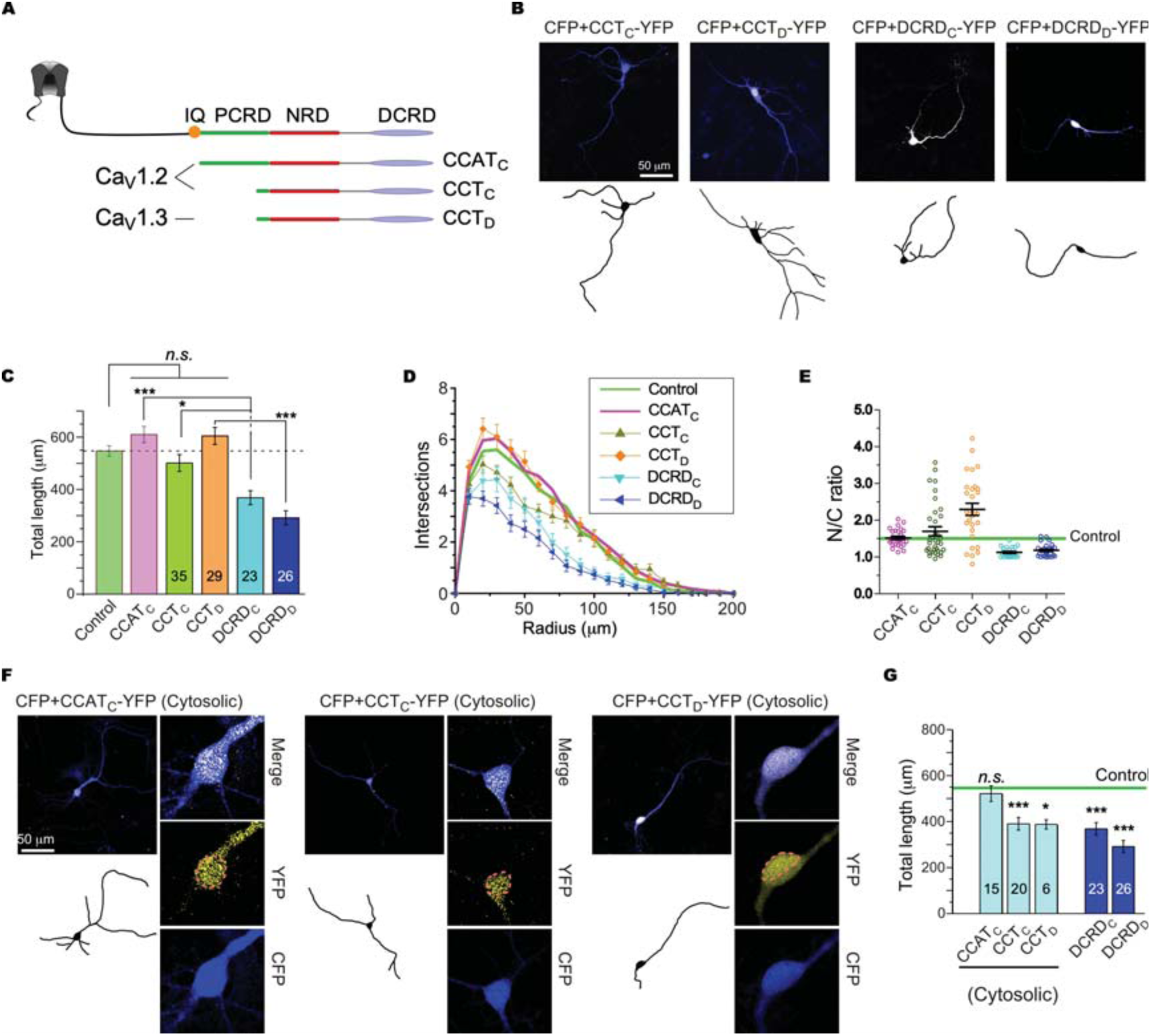
Intrinsic tendency of DCT peptides to downregulate neurite outgrowth in cortical neurons. (**A**) Illustration of three naturally occurring peptides encoded from DCTs of Ca_V_1.2 and Ca_V_1.3. CCAT_C_ contains the complete set of motifs including PCRD, NRD and DCRD. CCT_C_ and CCT_D_ are encoded by starting from the very end of PCRD all the way to the end of DCRD thus containing both NRD and DCRD. (**B**) Cortical neurons (DIV-7) expressing CFP and DCT variants tagged with YFP were imaged with fluorescence confocal microscopy. Representative confocal images (upper row, merged CFP and YFP channels) and neurite tracings (bottom row) are shown for CCT_C_, CCT_D_, DCRD_C_ and DCRD_D_, respectively. (**C** and **D**) Statistical summary of total neurite length (**C**) and *Sholl* analyses (**D**). Data for the Control and CCAT_C_ groups were adopted from Figure 1B,C. (**E**) Statistical summary of N/C ratio for all the 5 groups of DCT peptides. Horizontal line (green) indicates N/C ratio of GFP as the criteria of expression distributions to assign each neuron to either *nuclear* group (N/C ratio>1.5) or *cytosolic* group (N/C ratio<1.5). (**F**) Representative confocal images, neurite tracings and detailed cytonuclear distribution are shown for neurons with cytosolic CCAT_C_, CCT_C_ and CCT_D_, respectively. The envelop of the nucleus is highlighted by live-cell Hoechst 33342 staining (dotted circle in pink). (**G**) Total length for the neurons from the *cytosolic* groups: peptides of CCAT_C_, CCT_C_ and CCT_D_, and also the groups of DCRD_C_ and DCRD_D_ peptides. One-way ANOVA followed by Bonferroni or Dunnett for post hoc tests were used for (**C**) and (**G**), respectively (*, *p*<0.05; **, *p*<0.01; ***, *p*<0.001). Data were shown as means±SEM.

Whether and how these DCT variants of different size and compositions would exert CMI effects on native Ca_V_1 channels, e.g., neuronal Ca_V_1.3, have remained uncertain. Despite multiple evidence that DCT peptides inhibit Ca_V_1 channels (Hulme, Yarov-Yarovoy et al. 2006, Liu, Yang et al. 2017), no data thus far provide the direct proof of effectiveness for any DCT peptide on the full-length Ca_V_1 variants, e.g., Ca_V_1.3 containing its own set of PCRD and DCRD. Moreover, thus far it is still lacking direct evidence to support CMI as one inhibition modality for Ca_V_1-dependent transcriptional signaling and neurite outgrowth, with DCT being as either channel modules or standalone peptides.

Here, we first provided the basis to clarify the above questions and concerns. Notably, we unveiled that CMI potency of DCT peptides serves as mechanistic determinants for: 1) tunable channel inhibitor of Ca_V_1 and its Ca^2+^ influx; 2) tunable nuclear fraction of transcriptional regulators; and 3) tunable balance between the two signaling opponents (cytosolic/inhibitory *versus* nuclear/trophic signals of peptides) for neurite growth. In addition to its immediate significance to Ca_V_1 gating and signaling in neurons, a *de novo* scheme is emerging that one signaling molecule could autonomously tune the cyto-nuclear dynamics of its own, as the way to dynamically balance its opposed subcellular (cytosolic versus nuclear) roles of signaling, expected to broadly find its manifestations.

## Results

### Unexpected effects of DCT peptides on Ca_V_1 channels and neurons

As the earliest evidence regarding endogenous DCT peptides, CCAT_C_ has revealed its role as transcription factor to promote neurite outgrowth (Gomez-Ospina, Tsuruta et al. 2006). For validations under our experimental conditions, CCAT_C_ was transfected and overexpressed in DIV5 (5 days in vitro) cultured neurons from mouse cortex for two days (Figure 1A). Unexpectedly, CCAT_C_ did not cause any significant promotion on the length or complexity of neurites in contrast to previous reports(Gomez-Ospina, Tsuruta et al. 2006), even though there was seemingly a tendency of slight increase in neurite outgrowth (Figure 1B,C). Considering the potential CMI effects of DCT peptides (Hulme, Yarov-Yarovoy et al. 2006, Liu, Yang et al. 2017), such complications of CCAT_C_ might be due to the reduction of Ca_V_1-mediated Ca^2+^ influx and subsequent signaling (Wheeler, Groth et al. 2012, Li, Tadross et al. 2016), thus neutralizing its role as neurotrophic TF. However, to our surprise again, CMI of CCAT_C_ exhibited rather weak CMI on recombinant Ca_V_1.3 of full-length (α_1DL_), one major Ca_V_1 isoform expressed in cortical neurons (Striessnig, Pinggera et al. 2014), which is subject to competition gauged by DCT and apoCaM (Liu, Yang et al. 2010) (Figure 1D). In details, neither CDI nor VGA was altered, according to the respective index of CDI strength *S_Ca_* or peak Ca^2+^ current *I_Ca_* (Liu, Yang et al. 2017). CMI potency is defined by the normalized change (in percentage) of *S_Ca_* by comparing CCAT_C_ with the control. CMI potency of CCAT_C_ on α_1DL_ channels turned out to be rather low (Figure 1E), indeed inviting further investigations.

To resolve the above complications regarding the actual effects of DCT peptides, we systematically compared CCAT_C_, CCT_C_ and CCT_D_ containing intact NRD and DCRD domains from Ca_V_1.2 or Ca_V_1.3 (Figure 2A and Figure 2-figure supplement 1), and all these peptides are naturally present in native cells. In addition, DCRD_C_ and DCRD_D_ peptides encoded by the subset sequences of the CCT_C_/CCAT_C_ and CCT_D_ respectively were also examined side-by-side, serving as important controls due to their lack of NRD transactivation domain (Gomez-Ospina, Tsuruta et al. 2006, Dewenter, von der Lieth et al. 2017). Similar to CCAT_C_, CCT_C_ and CCT_D_ did not exhibit any significant effects (neither promotion nor retraction) on neurite length and complexity based on morphological analyses of confocal images. In clear contrast, DCRD_C_ and DCRD_D_ induced significant neurite retractions, presumably due to inhibitory effects on transcription and expression of neurotrophic genes (Figure 2B-D). Such destructive effects on neurites are supposedly the downstream of Ca_V_1 inhibition (CMI) by the peptides present in the cytosol (instead of the nucleus). Notably, the nucleus-cytosol ratio (N/C ratio) of DCT peptides appeared to be drastically different for the peptides with or without significant effects on neurite outgrowth. In details, the values of N/C ratio for peptides DCRD_C_ and DCRD_D_ in vast majority of neurons were smaller than the GFP control group (evenly distributed with N/C ratio ∼1.5) (Yang, Liu et al. 2018), confirming their dominance in cytosol; in contrast, the peptides CCAT_C_, CCT_C_ and CCT_D_ were more frequently found in the nucleus (N/C ratio>1.5) (Figure 2E). Moreover, by the criteria of N/C ratio at the cut-off value of 1.5, all the above neurons could be segregated into two (cytosolic versus nuclear) groups for each peptide, unmasking the tendency of cytosolic peptide (N/C ratio<1.5) to induce neurite retractions (Figure 2F,G). Similar to peptides DCRD_C_ and DCRD_D_, cytosolic CCT_C_ and CCT_D_ significantly attenuated neurite outgrowth. Cytosolic CCAT_C_ exhibited the tendency to inhibit Ca_V_1 signaling and consequent neurite outgrowth, while the overall effects of whole-cell CCAT_C_ seemed constructive (Figure 1B,C), both to confirm with further data.

### Tight correlations between peptide CMI and neurite retraction

Inhibition of Ca_V_1 channels is associated with downregulation of neuronal development (Nemzou N, Bulankina et al. 2006), supposedly by perturbations on neurotrophic signaling via Ca_V_1-mediated excitation-transcription coupling (Dolmetsch, Pajvani et al. 2001, Wong and Ghosh 2002, Wheeler, Groth et al. 2012, Li, Tadross et al. 2016). We hypothesized that the inhibitory profiles of cytosolic DCT peptides on neurite outgrowth might be due to their differential CMI. Based on the prior knowledge, the most critical DCT domain is the highly homologous DCRD among DCT variants (Figure 2-figure supplement 1). Except mysterious CCTA_C_, the other four peptides including DCRD_C_, CCT_C_, DCRD_D_ and CCT_D_ all clearly exerted CMI on recombinant full-length Ca_V_1.3 channels (in HEK293 cells), with the prominent features of concurrent attenuation on both CDI (*S_Ca_*) and VGA (*I_Ca_*) assuring the overall inhibition of Ca^2+^ influx (Liu, Yang et al. 2017) (Figure 3A), also indexed by the primary index of CMI potency (Figure 3B). Before this work, it was uncertain whether DCT peptides would effectively act on the full-length (native-form) Ca_V_1 to induce channel inhibition. Here, we confirmed the effectiveness of peptide-form CMI. Notably, we relieved the uncertainty regarding the peptide of DCRD_C_, which was not supposed to induce CMI considering the apparent ineffectiveness of CCAT_C_ peptide (on Ca_V_1.3) and DCRD_C_ domain (on Ca_V_1.2) (Hulme, Yarov-Yarovoy et al. 2006). Further mutagenesis trimmed down on DCRD_F_ to the core domain for CMI: the segment containing residues from 17 to 66 of high homology across Ca_V_1 family especially for Ca_V_1.2-1.4 (Figure 3-figure supplement 1), consistent with potent CMI of DCRD_C_, DCRD_D_ and DCRD_F_. Quantitatively, CMI potency and neurite length were inversely correlated for DCT peptide variants (Figure 3C), providing the direct evidence that cytosolic DCT peptides attenuate neurite outgrowth in accordance to inhibition of Ca^2+^ influx through the channel, potentially by downregulation of excitation-transcription coupling mediated by cortical Ca_V_1 channels.

**Figure 3.**
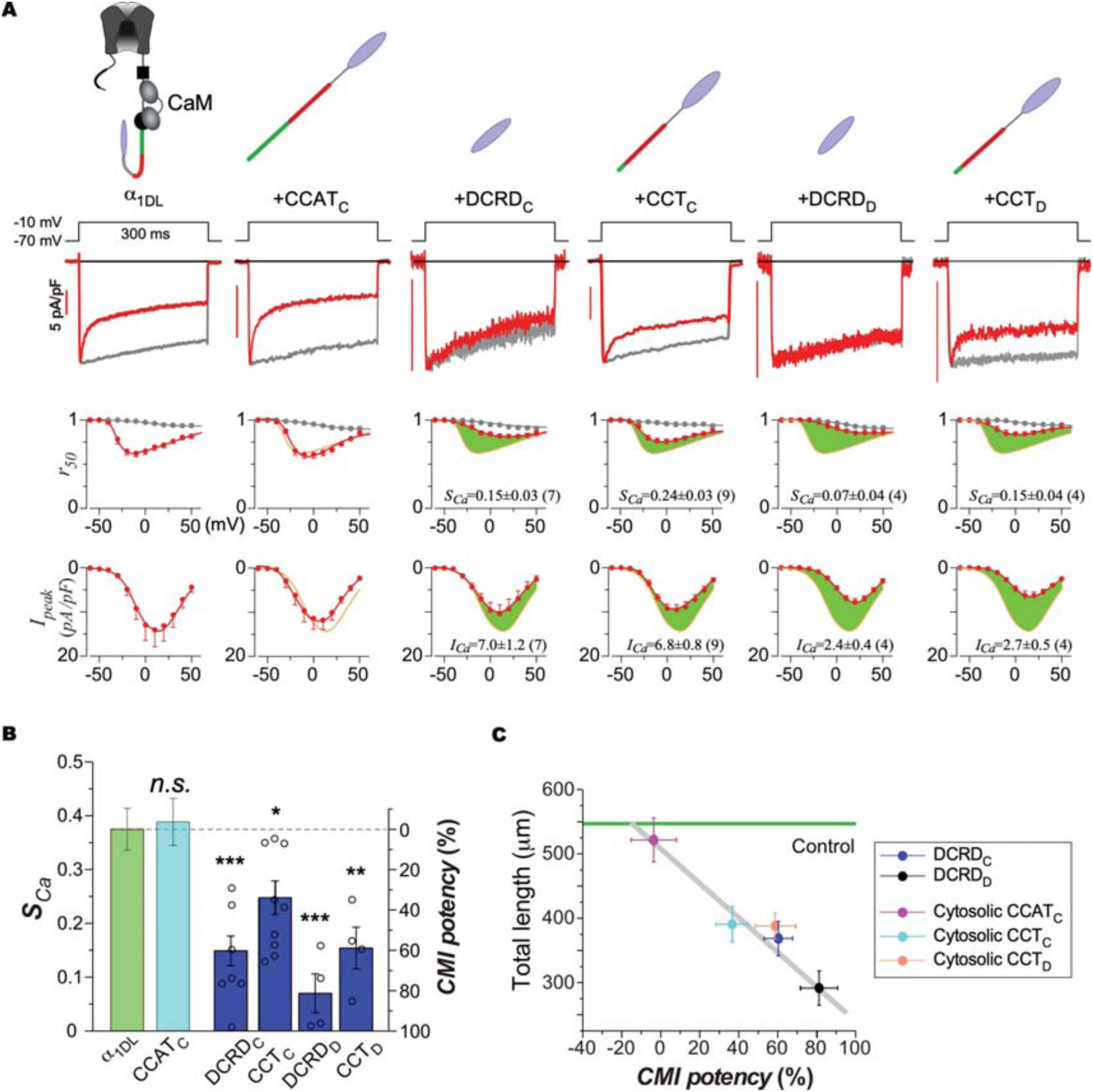
Distinct CMI potency of DCT peptides on Ca_V_1.3 and its correlations with neurite outgrowth. (**A**) CMI potency across DCT peptide variants. DCT peptides (cartoon on the top), representative current trace (first row), profiles to evaluate activation (*S_Ca_*) and inactivation (*I_Ca_)* are shown in order. Green areas are to highlight the differences in activation and inactivation profiles between control and peptide groups. Data of α_1DL_ control and CCAT_C_ are adopted from Figure 1D for comparison. (**B**) Statistical summary of *S_Ca_* and corresponding CMI potency. (**C**) Correlations between CMI potency and total length for neurons with peptide expression in the cytosol. Data of neurite length are adopted from Figure 2G. One-way ANOVA followed by Dunnett for post hoc tests were used for (**B**) (*, *p*<0.05; **, *p*<0.01; ***, *p*<0.001). Data were shown as means±SEM.

### Contributions of PCRD and DCRD to differential CMI of DCT variants

CCAT_C_ remained mysteriously ineffective (Figure 1 and Figure 3). In contrast to CCAT_C_, shorter peptides of DCRD_C_ and CCT_C_ have strong CMI, suggesting some unknown auto-inhibitory factor might affect CCAT_C_. Moreover, full-length Ca_V_1.2 does not show significant difference in CDI with or without DCT (Peterson, DeMaria et al. 1999, Hulme, Yarov-Yarovoy et al. 2006), also inconsistent with capability of channel inhibition intrinsic to DCRD_C_ domain. We again performed systematic analysis over DCT variants to seek the hint of potential resolution. Firstly, by 3-cube FRET (Förster resonance energy transfer) imaging as 2-hydrid binding assays, the capabilities of DCRD peptides to bind channel upstream motifs (thus exerting CMI) were quantified by *FR-D_free_* curves (Figure 4A-C). Effective dissociation equilibrium constant *K_d_* (units in donor-cube fluorescence intensity) is used to quantify the binding affinity, and smaller values of *K_d_* indicate stronger interactions. The FRET pairs of CFP-tagged DCRD_X_ peptides (X=S, D, C and F representing Ca_V_1.1-1.4) and YFP-tagged preIQ_3_-IQ_D_-PCRD_D_ exhibited a series of curves fitted by a gradient of *K_d_* values (Figure 4A), among which DCRD_F_ from Ca_V_1.4 exhibited the strongest binding affinity (*K_d_*=1.7×10^3^), followed by DCRD_D_ (*K_d_*=3.5×10^3^), DCRD_C_ (*K_d_*=7.8×10^3^) and DCRD_S_ (*K_d_*=8.6×10^3^). Similarly, the relative capabilities of PCRD_X_ (X=S, D, C and F) in CMI were examined in the context of YFP-preIQ_3_-IQ_D_-PCRD_X_ and CFP-DCRD_F_. The results revealed a similar order in binding affinities (starting from the strongest): PCRD_F_, PCRD_D_, PCRD_C_ and PCRD_S_ (Figure 4B). The difference in *K_d_* between PCRD_C_ (*K_d_*=5.9×10^3^) and PCRD_D_ (*K_d_*=1.8×10^3^) is even more dramatic than that between DCRD_C_ and DCRD_D_ (3.3-fold versus 2.1-fold), suggesting that the rather weak CMI for DCT_C_ or Ca_V_1.2 is mainly due to PCRD_C_. To summarize and compare, indexed by *K_d_* of the pair of DCRD_F_ and preIQ_3_-IQ_D_-PCRD_D_ as the reference (noted as PCRD_D_/DCRD_F_ or P_D_/D_F_), all PCRD_X_/DCRD_X_ (or P_X_/D_X_) combinations were evaluated as relative *K_d_* (P_X_/D_X_) normalized by *K_d_* (P_D_/D_F_) (Figure 4C). Besides the experimental values (indicated as exp. in Figure 4C), the relative *K_d_* values can be estimated for other combinations without being actually measured here by FRET. For P_C_/D_C_ (Ca_V_1.2) and P_S_/D_S_ (Ca_V_1.1), the actual results (11.2 and 18.7) were also obtained from experiments (Figure 4-figure supplement 1A,B), proved to be highly consistent with the predicted *K_d_* values (13.1 and 18.5), as additional validations of Figure 4C. In this context, the three natural peptides of CCT_D_, CCT_C_ and CCAT_C_ in principle could be represented by P_D_/D_D_, P_D_/D_C_ and P_C_/D_C_ respectively, in agreement with the order of corresponding CMI potency (Figure 3).

**Figure 4.**
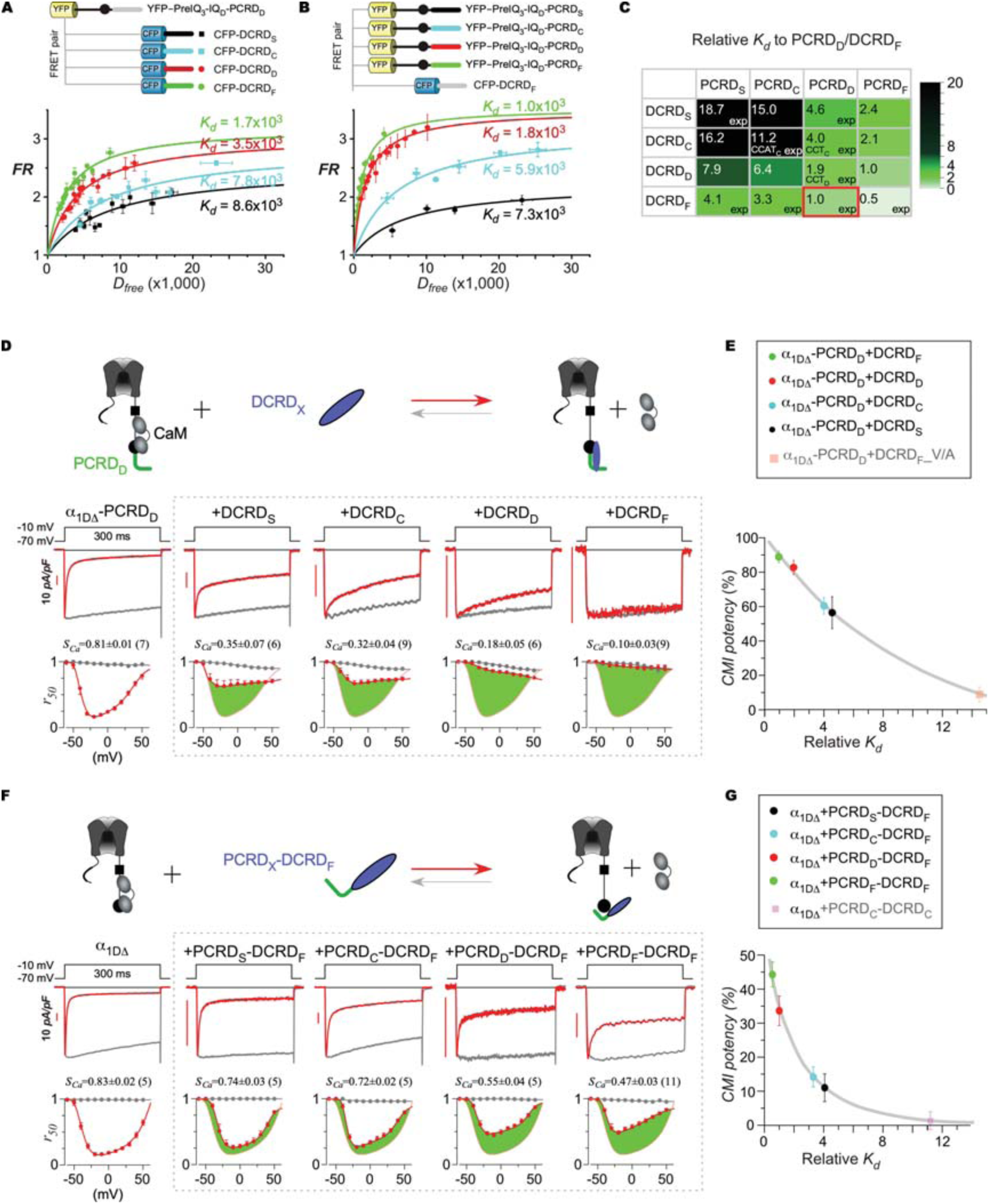
Quantitative analyses on PCRD and DCRD variants unveil CMI determinants. (**A**) Binding curves between the channel and DCRD peptides were quantified by 2-hydrid 3-cube FRET, for each pair between CFP-DCRD_X_ from Ca_V_1 (X=S, D, C and F) and YFP-preIQ_3_-IQ_D_-PCRD_D_. *D_free_* and *FR* represent free donor concentration and FRET ratio, respectively. The binding affinity *K_d_* for each binding curve was achieved by iterative fitting as the major index to compare the relative strength among the interactions. (**B**) Similar to the measurements for DCRD variants in (**A**), PCRD_X_ peptides across Ca_V_1 family (X=S, D, C and F) were also quantified by FRET for the interactions between YFP-preIQ_3_-IQ_D_-PCRD_X_ and CFP-DCRD_F_. (**C**) Summary of relative *K_d_* values for PCRD_X_-DCRD_X_ peptide variants. The *K_d_* of the binding between DCRD_F_ and preIQ_3_-IQ_D_-PCRD_D_ was set as the reference, based on which the relative *K_d_* of all PCRD/DCRD combinations were calculated as the ratio of *K_d_* (PCRD_X_/DCRD_X_) to *K_d_* (PCRD_D_/DCRD_F_). Experimental values were marked as “exp”, while the values of other combinations were predicted by extrapolations. The strength of relative affinity (*K_d_*) was indicated by a gradient of colors (color bar, right). The three native forms of DCT peptides are also labelled as CCAT_C_, CCT_C_, and CCT_D_, correspondingly. (**D**) Comparison of CMI potency among DCRD_X_ peptides. As in the cartoon illustration of CMI scheme (top), DCRD_X_ could coordinate with IQ_D_ and PCRD_D_ motifs on α_1DΔ_-PCRD_D_ to compete apoCaM off the channel. Representative Ca^2+^ traces and inactivation profiles are for α_1DΔ_-PCRD_D_ alone or with different DCRD_X_ isoforms. (**E**) Relationship between relative *K_d_* and CMI potency for DCRD_X_ peptides. Besides the four peptides directly from Ca_V_1.1-Ca_V_1.4, one additional peptide was the mutant DCRD_F__V/A (Figure 4-figure supplement 2). The smooth line connecting the five points (peptides) is for visual guidance of the tuning trend. (**F**) Comparison of CMI potency among PCRD_X_-DCRD_F_ peptides. As in the cartoon illustration of CMI scheme (top), PCRD_X_-DCRD_F_ could coordinate with IQ_D_ on α_1DΔ_ to compete apoCaM off the channel. Representative Ca^2+^ traces and inactivation profiles are for α_1DΔ_ alone or with different PCRD_X_-DCRD_F_ isoforms. (**G**) Relationship between relative *K_d_* and CMI potency for PCRD_X_-DCRD_F_ peptides. Besides the four peptides of PCRD_X_-PCRD_F_, one additional peptide was PCRD_C_-DCRD_C_ (Figure 4-figure supplement 1). The smooth line connecting the five points (peptides) is to visually guide the tuning trend. Data were shown as means±SEM.

Besides above evidence of the linkage between binding affinity and peptide CMI, electrophysiological analyses were conducted to systematically examine the effects of different peptide variants on channel gating. Firstly, DCRD_X_ peptides were evaluated in the context of recombinant α_1DΔ_-PCRD_D_ channels, which lack the critical DCRD domain thus leading to ultra-strong CDI with ample dynamic range for potential modulations. All of the four DCRD_X_ peptides (encoding DCTs of Ca_V_1.1-1.4) exerted CMI of different potency on α_1DΔ_-PCRD_D_ channels, directly reflected into their CDI profiles (Figure 4D). Most importantly, a tight correlation was unveiled between CMI potency and relative *K_d_* for these DCRD peptides (Figure 4E). Furthermore, DCRD_F__V/A peptide (with the critical mutation V41A) strongly attenuated both peptide binding and CMI potency(Liu, Yang et al. 2010) (also see Figure 4-figure supplement 2), nicely falling onto the tuning curve of *K_d_*-CMI. Likewise, the role of PCRD in CMI was also examined in the context of peptides PCRD_X_-DCRD_F_ (i.e., P_X_-D_F_) co-expressed with α_1DΔ_ channels. P_S_-D_F_ and P_C_-D_F_ peptides exhibited rather weak inhibition, much less potent than P_D_-D_F_ and P_F_-D_F_ peptides that significantly attenuated CDI of α_1DΔ_ channels (Figure 4F). Overall, the four peptides of PCRD_X_-DCRD_F_ exhibited a similar but quantitatively different tuning curve of CMI potency, indexed by relative *K_d_* varying with each *P_X_* (Figure 4G). CMI tuning curves of PCRD and DCRD (Figure 4E,G) together as the molecular determinants account for the differential CMI strength of DCT peptides or motifs across Ca_V_1.1 to Ca_V_1.4 (Hulme, Yarov-Yarovoy et al. 2006, Stroffekova 2008, Liu, Yang et al. 2010, Ohrtman, Romberg et al. 2015). In addition, the weak *K_d_* and CMI of PCRD_C_-DCRD_C_ (Figure 4-figure supplement 1B,C) in agreement with the tuning curve suggested its potential applicability to P_X_-D_X_ peptides (Figure 4G). We here discovered that PCRD acts as one major factor to determine overall CMI of DCT. The ultra-weak CMI potency of DCT_S_ and DCT_C_ should be largely attributed to PCRD, in comparison with DCRD that has relatively less contribution.

### Mechanisms of weak CMI of peptide CCAT_C_

Although the weak PCRD_C_ has been unveiled in the above settings, it is still unclear why full-length Ca_V_1.3 which contains P_D_ capable of strong CMI is also merely regulated by CCAT_C_. To dissect out the exact mechanisms, we examined the CMI potency of peptide CCAT_C_ on α_1DΔ_-PCRD_D_ channels, which can be decomposed into two possible combinations or components (Figure 5A,C). The first component (I) represents the inter-molecular combination of P_D_ from the channel and D_C_ from the peptide, exhibiting CMI of substantial strength (Figure 4D). The second component (II) represents the intra-molecular combination of P_C_ and D_C_ both from the peptide DCT_C_ (equivalent to CCAT_C_), which has rather weak CMI (Figure 4-figure supplement 1). As expected, the compound effects of CCAT_C_ on α_1DΔ_-PCRD_D_ resulted into modest CMI potency falling in between the two decompositions I and II, demonstrating the similar principle as in Figure 4 that PCRD and DCRD synergistically determine CMI potency of the peptides, e.g., CCAT_C_ where the weak PCRD_C_ plays a dominant role (Figure 5C). Engineered peptide P_C_-D_F_ further consolidated the above insights (Figure 5B,D). Similar to P_C_-D_C_ or CCAT_C_, the effects of P_C_-D_F_ on α_1DΔ_-P_D_ could also be decomposed as intra- and inter-molecular combinations (Figure 4D,F), also resulting into intermediate CMI potency.

**Figure 5.**
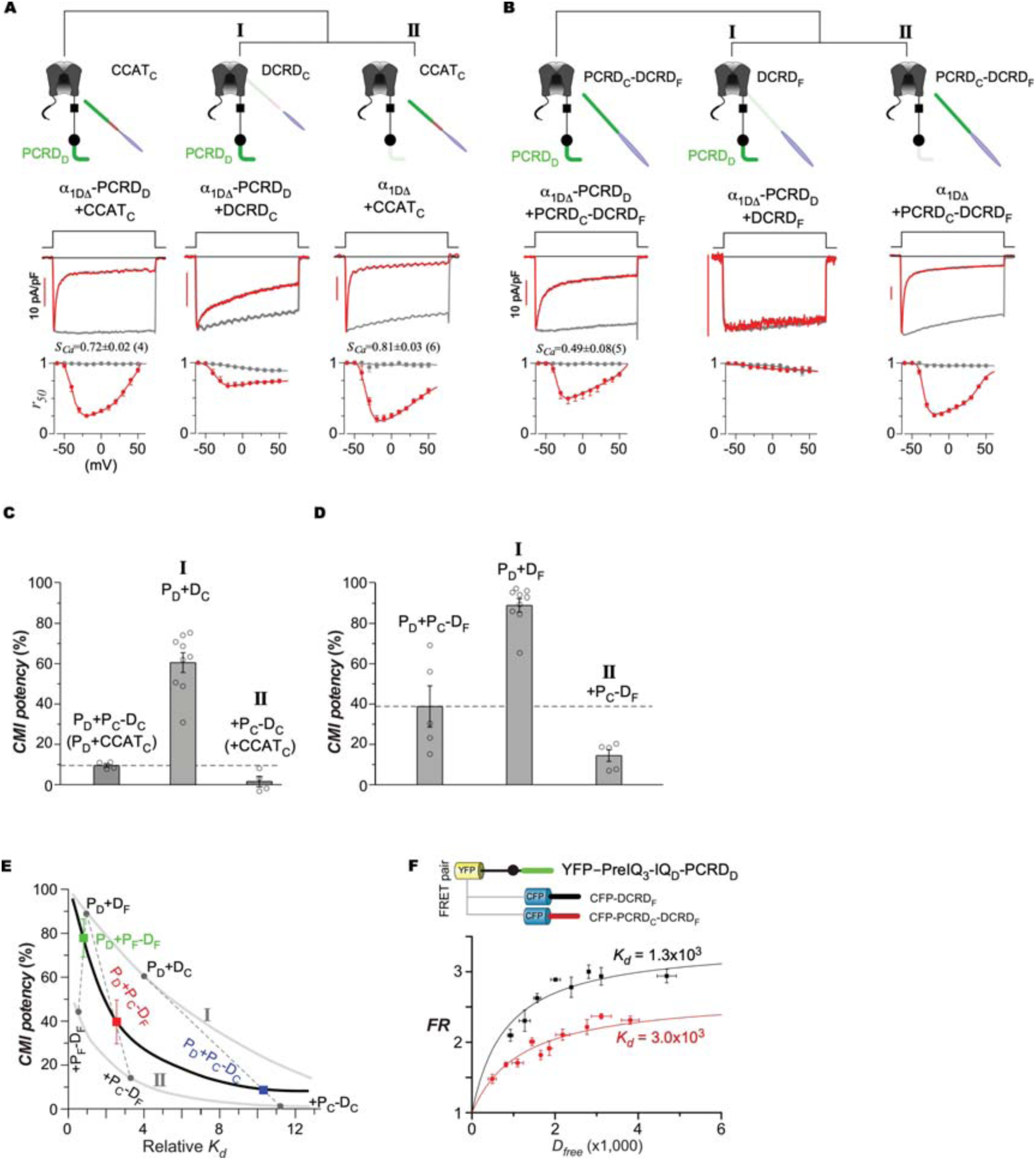
Compound CMI effects account for weak CCAT_C_ modulation of Ca_V_1.3. (**A**) Decomposition of CCAT_C_ effects on α_1DΔ_-PCRD_D_ channels. CMI effects of CCAT_C_ could be divided into two components (top row). The first component (I) represents CMI of DCRD_C_ (contained in CCAT_C_) on α_1DΔ_-PCRD_D_, where PCRD and DCRD motifs required by CMI are contributed by two individual molecules (inter-molecular combination): α_1DΔ_-PCRD_D_ and CCAT_C_, respectively. Data of component I was adopted from Figure 4D; whereas data of component II are almost identical to Figure 4-figure supplement 1 due to minimum difference between CCAT_C_ and PCRD_C_-DCRD_C_ in terms of CMI. The second component (II) represents CMI of CCAT_C_ on α_1DΔ_, where PCRD and DCRD motifs required by CMI are from the same molecule (intra-molecular combination): both from CCAT_C_. Representative Ca^2+^ traces (middle row) and inactivation (bottom row) profiles are shown in each column for the effects of CCAT_C_ and two different combinations. (**B**) Decomposition of peptide PCRD_C_-DCRD_F_ effects on α_1DΔ_-PCRD_D_ channels. Similar to (**A**), CMI effects of the DCT peptide are decomposed into two components by DCRD_C_ on α_1DΔ_-PCRD_D_ (I, inter-molecular combination) and by PCRDC-DCRDF on α_1DΔ_ (II, intra-molecular combination) (top row). Data of component I and II were adopted from Figure 4D and Figure 4F, respectively. (**C** and **D**) Statistical summary of CMI potency for the peptides of CCAT_C_ and PCRD_C_-DCRD_F_ and the two components for each peptide. The dashed lines were drawn to compare the relative CMI potency of the three groups for CCAT_C_ or PCRD_C_-DCRD_F_ effects. The compound effects of α_1DΔ_-PCRD_D_+PCRD-DCRD (P_D_+P-D), can be decomposed into α_1DΔ_-PCRD_D_+DCRD (component I, P_D_+D) and α_1DΔ_+PCRD-DCRD (component II, +P-D). (**E**) Relationship between CMI potency and relative *K_d_* for the effects of PCRD-DCRD peptides on α_1DΔ_-PCRD channels. The gray lines at the top (I) and bottom (II) are adopted from Figure 4E and Figure 4G, respectively. For CCAT_C_, its compound CMI effects are nearly identical to P_D_+P_C_-D_C_ (blue, **C**), based on which the equivalent *K_d_* (relative) can be estimated by connecting the two data points of components I and II on the respective gray lines. Similarly, compound effects of P_D_+P_C_-D_F_ (red, **D**) and P_D_+P_F_-D_F_ (green, Figure 5-figure supplement 1) were also attributed to the corresponding data points. The black line linking the three (blue, red and green) data points of compound effects is to provide a visual guidance of the turning trend. (**F**) Measurement of *K_d_* for the interaction between preIQ_3_-IQ_D_-PCRD_D_ and PCRD_C_-DCRD_F_, in comparison with the control group of DCRD_F_, by 2-hybird 3-cube FRET. Data were shown as means±SEM.

Systematically, the quantitative relationships between peptide (binding) and its function (CMI) could be achieved (for different peptide variants) by correlating the compound effect of long peptide with its two (inter- and intra-molecular) combinations (Figure 5E). In the context of P_X_-D_X_ peptides and α_1DΔ_-P_D_ channels, the two curves for components I and II, which represent inter- and intra-molecular combinations, are essentially identical to the curves of D_X_ peptides on α_1DΔ_-P_D_ (Figure 4E) and of P_X_-D_X_ peptides on α_1DΔ_ (Figure 4G), respectively. For P_C_-D_F_ peptide on α_1DΔ_-P_D_, its compound CMI potency should be between the two points of decomposed I (α_1DΔ_-P_D_+D_F_) and II (α_1DΔ_+P_C_-D_F_), presumably falling onto the line that connects the two points (I and II) (marked by the red point in Figure 5E and Figure 5B). As validation, *K_d_* (relative *K_d_* =2.4) estimated from above procedure approximates the experimental value by FRET between P_C_-D_F_ and preIQ_3_-IQ_D_-P_D_ (relative *K_d_* =2.3, Figure 5F). Similarly, compound effects on α_1DΔ_-P_D_ and equivalent *K_d_* of P_F_-D_F_ (green point in Figure 5E and Figure 5-figure supplement 1) and P_C_-D_C_ (blue point in Figure 5E and Figure 5A) could also be decomposed and predicted, respectively. The middle curve corresponding to the compound CMI effects on full-length channel, together with the other two curves, proposes a useful platform for systematic analysis of DCT peptides, including those not included in this study: CCT_S_ (Hulme, Konoki et al. 2005), CCAT_D_, CCT_F_ and CCAT_F_. Finally, the difference in binding affinity between D_F_ and P_C_-D_F_ was unveiled by FRET, providing further evidence that the weak CMI of peptide CCAT_C_ is mainly attributed to its weak P_C_ that essentially interferes with DCT/CMI endogenous to the native channel.

### Cyto-nuclear dependence of peptide signaling reconciles its opposing effects

The data thus far unveiled that cytosolic DCT peptides negatively regulate dendritogenesis, in a variant-dependent manner intrinsically tuned by the CMI strength. Collectively, the apparent contradiction of peptide effects could simply reflect the differential roles of peptides in the cytosol as opposed to in the nucleus. To validate the potential scheme of opposing cytonuclear signaling, we first revisited CCT_D_ which widely distributed across the whole cell, indicated by N/C ratio values of wide range (Figure 2E). Significant effects of dual directions were evident: for nuclear or CCT_D_ (N) group (N/C ratio>1.5) neurite outgrowth was dramatically promoted; whereas neurons in CCT_D_ (C) group (N/C ratio<1.5) exhibited neurite retraction similar to DCRD_D_ group (Figure 6A-C). To more explicitly examine CCAT_C_ and CCT_C_, short tags of nuclear export signal (NES) and nuclear localization signal (NLS) were fused to the N-terminus of the peptide. NES-tagged CCT_C_ and CCAT_C_ were clearly located in the cytosol and exhibited differential levels of neurite retraction, consistent with the tuning trend shown earlier (Figure 3). In contrast, NLS-tagged CCAT_C_ and CCT_C_ were constrained in the nucleus, acting as morphogenic factors to facilitate neurite outgrowth, as expected from previous reports(Gomez-Ospina, Tsuruta et al. 2006) (Figure 6D-G). When specifically overexpressed in the nucleus, peptides of CCAT_C_, CCT_C_ and CCT_D_ appeared to promote neurite outgrowth to comparable levels, suggesting their similar roles as nuclear TFs (Figure 6H), in contrast to downregulatory roles of these peptides when present in the cytosol (Figure 2G), the latter of which is also subject to variant-, affinity- or CMI-dependent tuning.

**Figure 6.**
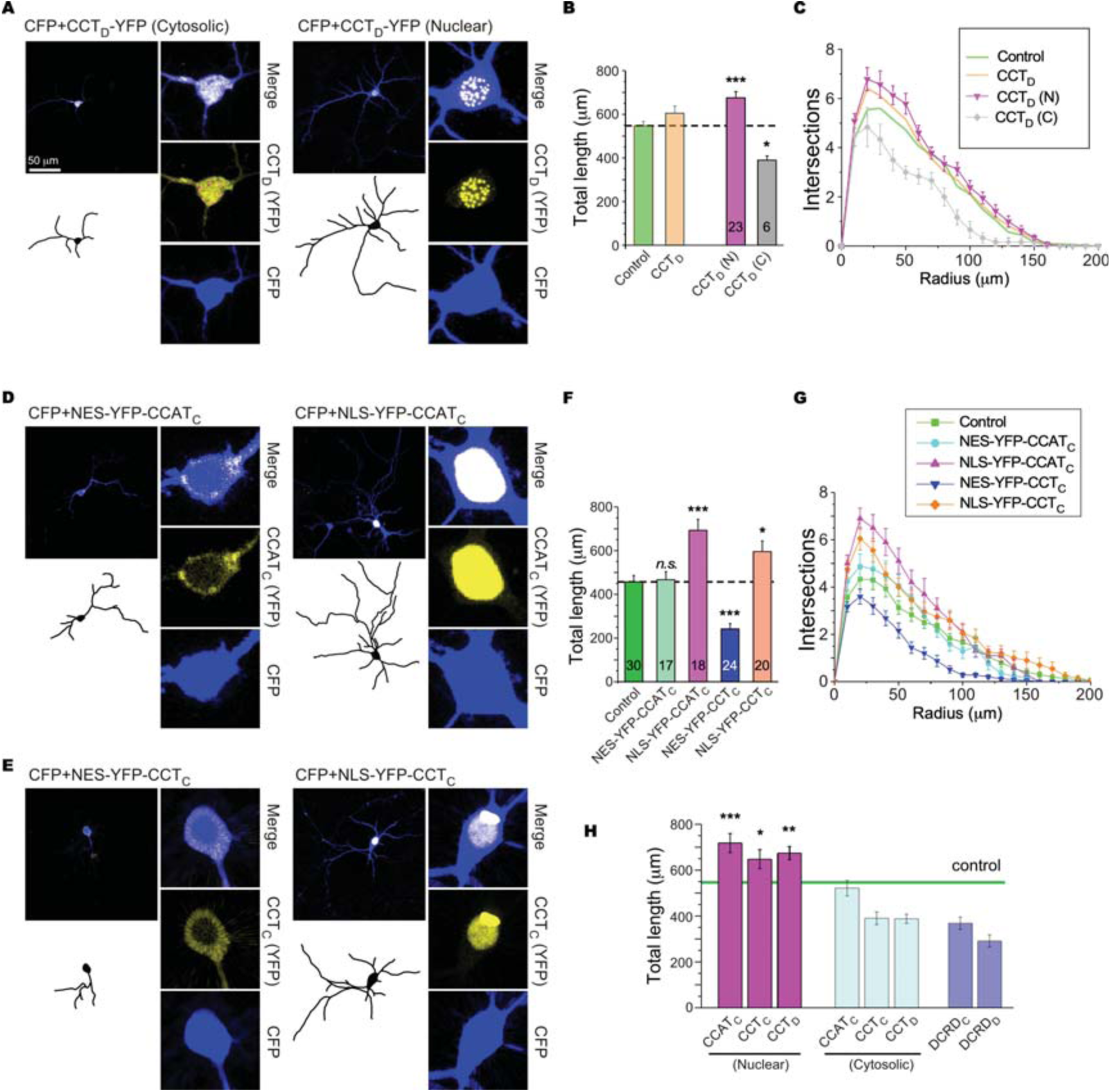
Cyto-nuclear dependence reconciles the opposing effects of DCT peptides. (**A**) Cortical neurons with CCT_D_ fragments were grouped into two categories: nuclear *versus* cytosolic, i.e., CCT_D_ (N) and CCT_D_ (C) by the same criteria of N/C ratio as in Figure 2 (CCT_D_ fluorescence with N/C ratio>1.5 for the nuclear group; and N/C ratio<1.5 for the cytosolic group). (**B** and **C**) Total neurite length (**B**) and *Sholl* analyses (**C**) for the groups of Control, total CCT_D_, CCT_D_ (C), CCT_D_ (N) and DCRD_D_. Original data for the control and total CCT_D_ groups were adopted from Figure 2C,D. (**D** and **E**) NES and NLS were fused to N-terminus of YFP-CCAT_C_ or YFP-CCT_C_ to constrain the distribution of DCT peptides in either the cytosol or the nucleus, respectively. (**F** and **G**) Total neurite length (**F**) and *Sholl* analyses (**G**) for the groups of Control, NES-YFP-CCAT_C_ versus NLS-YFP-CCAT_C_, and NES-YFP-CCT_C_ versus NLS-YFP-CCT_C_. (**H**) Total neurite lengths for the neurons overexpressing CCAT_C_, CCT_C_ and CCT_D_ peptides localized in the nucleus versus the cytosol, as compared with DCRD_C_ and DCRD_D_ (latter data were adopted from Figure 2G). One-way ANOVA followed by Dunnett for post hoc tests were used for (**B**), (**F**) and (**H**) (*, *p*<0.05; **, *p*<0.01; ***, *p*<0.001). Data were shown as means±SEM.

### Cytonuclear translocation in response to Ca^2+^ influx via Ca_V_1 is regulated by CMI

Such opposing effects could be largely cancelled out by each other, thus responsible for the observed ambiguity of overall peptide effects on neurite outgrowth. However, it still remained puzzling how such counteraction/balance could be delicately achieved for all these naturally occurring peptides since one side of the balance is DCT (N) with nearly identical capability to promote dendritogenesis; whereas DCT (C) on the other side could have drastically different CMI and thus different attenuation of neurite outgrowth. First, the spatial distribution of DCT peptides was examined in the context of neurite outgrowth. As depicted by the scatter plots to correlate N/C ratio with neurite length, CCAT_C_, CCT_C_ and CCT_D_ were all spatially dynamic spanning across the cyto-nuclear space in the whole-cell (Figure 7A-C), in contrast to DCRD_C_ and DCRD_D_ constrained within the cytosol (thus exclusively acting as inhibitors of dendritogenesis) (Figure 7-figure supplement 1). Even for the three natural peptides, the spatial patterns of their distributions were quantitatively different, as indexed by the dynamic range between minimum and maximum N/C ratio values. CCAT_C_ exhibited rather narrow N/C ratio range or least presence in the nucleus. In contrast, CCT_D_ was much more spatially dynamic with clear preference of nuclear localization. The dynamic range of cytonuclear distribution seemingly corresponds to the gradient of CMI potency in the order of CCAT_C_, CCT_C_ and CCT_D_ (from weak to strong, Figure 2E). All these data support that Ca_V_1 and its specific Ca^2+^ influx should play exceptional roles in the behaviors and effects of the peptides in cortical neurons.

**Figure 7.**
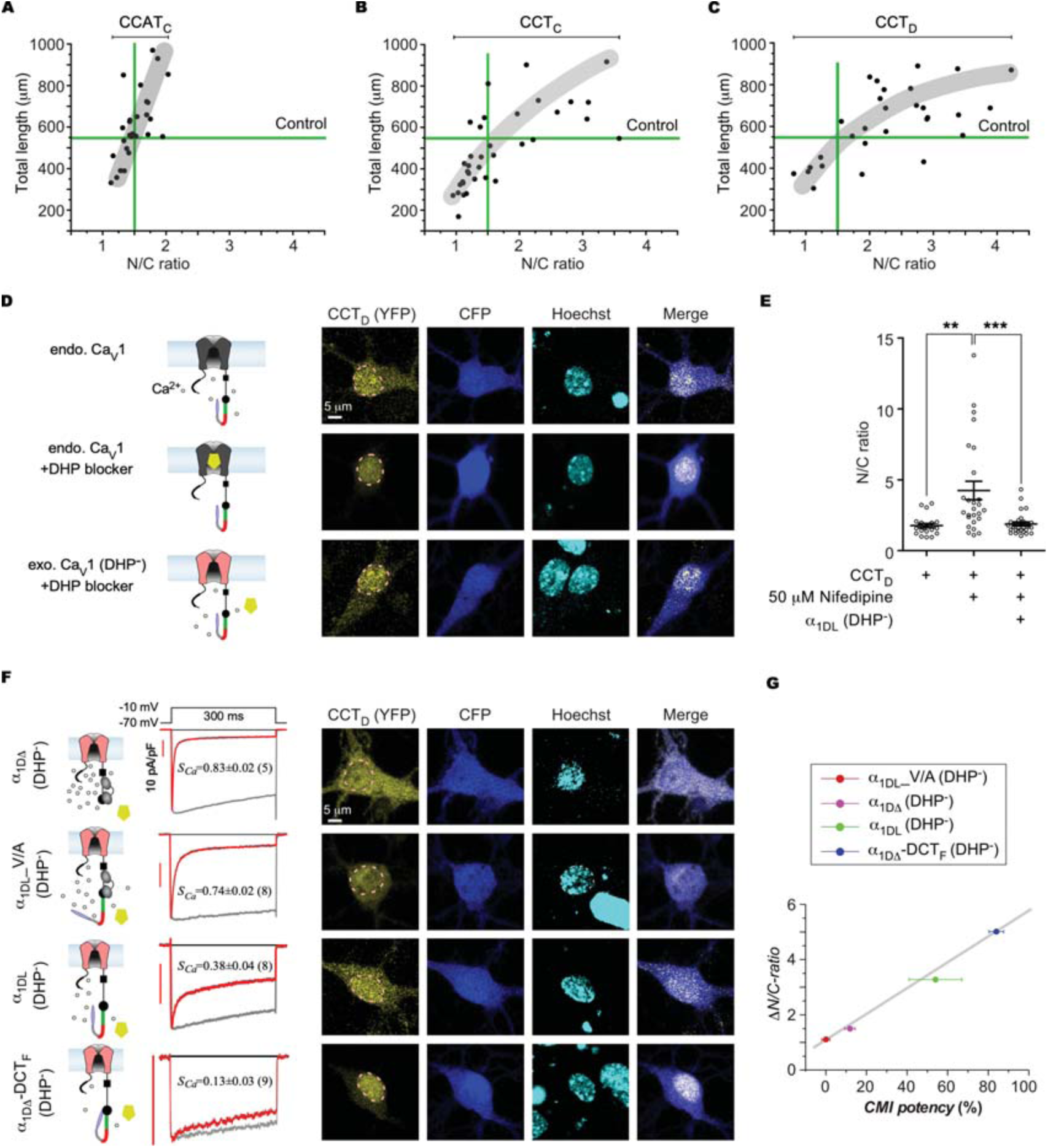
Cyto-nuclear distribution of DCT Peptides is tightly regulated by Ca^2+^/Ca_V_1. (**A-C**) Cyto-nuclear distribution (indexed by N/C ratio) of DCT peptides in correlation with the total neurite length. Horizontal and vertical lines in green represent the Control group (GFP expressed in cortical neurons). The range of N/C ratio was indicated at the top for each peptide as the index of its spatial (cyto-nuclear) dynamics. The thick lines in grey are to visually guide the trend of the correlation between the total neurite length and N/C ratio. (**D** and **E**) Effects of dihydropyridine (DHP) on cortical neurons expressing endogenous and/or exogenous Ca_V_1 channels together with CCT_D_ peptides. As shown in the cartoon (**D**, left), endogenous Ca_V_1 channels in cortical neurons mediate Ca^2+^ influx (upper); DHP (50 μM nifedipine) would specifically block the Ca_V_1 channels thus reducing the Ca^2+^ influx (middle); and overexpression of Ca_V_1 (DHP-) channels (mutant Ca_V_1.3 insensitive to DHP) could rescue Ca^2+^ influx while DHP blocks endogenous Ca_V_1 currents (bottom). Representative confocal images of YFP (CCT_D_ distribution), CFP (soma contour), Hoechst (nuclear envelop) and YFP/CFP merge for the three experimental groups (**D**, right), for which the values of N/C ratio were illustrated for individual neurons with the statistical summary (**E**). (**F**) Ca_V_1 variants with different CMI potency and the cyto-nuclear distribution of CCT_D_. Cortical neurons were co-expressed with CCT_D_ with one of the four DHP-resistant Ca_V_1.3 variants: α_1DΔ_ (lacking DCT), α_1DL__V/A (with the key mutation of V to A at DCRD), α_1DL_ (control), and α_1DΔ_-DCT_F_ (chimera with ultra-strong DCT), with representative current traces shown on the right side of the cartoon illustrations. Cortical neurons were treated with 50 μM nifedipine to block endogenous Ca_V_1 channels but spare the exogenous DHP^-^ channels, for which confocal images depicted the differential cyto-nuclear localizations of CCT_D_ peptides, in the similar fashion as in (**D**). (**G**) Close correlation between CMI potency and spatial dynamics for CCT_D_ peptides in cortical neurons for the four different variants of Ca_V_1.3 channels. The dynamic cyto-nuclear distribution of the peptide in individual neurons was indexed by the overall range in the values of N/C ratio (*ΔN/C-ratio*). One-way ANOVA followed by Bonferroni for post hoc tests were used for (**E**) (**, *p*<0.01; ***, *p*<0.001). Data were shown as means±SEM.

In fact, previous studies have demonstrated that the mobility of DCT peptides from nucleus to cytosol is regulated by intracellular Ca^2+^ rise(Gomez-Ospina, Tsuruta et al. 2006, Lu, Sirish et al. 2015), where Ca^2+^ influx through Ca_V_1 channels makes significant contributions(Gomez-Ospina, Tsuruta et al. 2006), as confirmed in our experiments (Figure 7-figure supplement 2). When Ca_V_1 activities at the basal conditions (5 mM [K^+^]_o_) were blocked by 50 μM nifedipine (DHP derivatives), CCT_D_ exhibited an increasing tendency of nuclear localization. Moreover, N/C ratio of CCT_D_ could be reverted back to the control level when overexpressing dihydropyridine insensitive channels (α_1DL_ DHP^-^) (Figure 7D,E), supporting the unique importance of Ca_V_1 in CCT_D_ translocation. In the context of Ca_V_1 DHP^-^ channels and DHP blockage of endogenous Ca_V_1 channels, we further investigated the effects of Ca_V_1 inhibition (CMI) on peptide distribution. Each of the four kinds of DHP-resistant Ca_V_1.3 variants, α_1DΔ_, α_1DL__V/A, α_1DL_ or α_1DΔ_-DCT_F_, in the order of increasing CMI potency (endogenous) and thus decreasing Ca^2+^ influx, was examined in cortical neurons overexpressing CCT_D_ in 50 μM nifedipine (Figure 7F). As expected, CCT_D_ in neurons of α_1DΔ_-DCT_F_ (DHP^-^) group was constricted in the nucleus and exhibited the largest dynamic range of N/C ratio values, corresponding to the strongest CMI potency intrinsic to the channels. In contrast, in neurons expressing α_1DΔ_ DHP^-^ channels that have the weakest CMI (due to lacking DCT), CCT_D_ was more constrained within the narrowest band of N/C ratio. Collectively, the four α_1D_ variants altogether unveiled a tight correlation between Ca^2+^ influx (gauged by CMI) and peptide (cytonuclear) distribution in cortical neurons (Figure 7G).

On the other hand, different peptides varying with the level of CMI should also similarly regulate their own spatial dynamics. To examine the potential correlation between peptide’s CMI and spatial dynamics, we took advantage of the critical mutant CCT_D_ carrying single V/A mutation in DCRD domain that drastically reduced CMI acting on α_1DL_ channels (Figure 8A-C). Subsequent transfection of CCT_D__V/A peptide into cortical neurons unveiled that in comparison with WT CCT_D_ the spatial dynamics of CCT_D__V/A was effectively attenuated (Figure 8D,E). Meanwhile, clear difference in neurite outgrowth was also evidenced from the mutant peptide which nearly lost its inhibitory effects when present in the cytosol (Figure 8F,G). Notably, V/A mutation did not affect the facilitory effects of nuclear CCT_D_ on neurite outgrowth. In summary, the tight correlations between CMI potency and N/C-ratio range for CCAT_C_, CCT_C_, CCT_D_ and also CCT_D__V/A peptides support the notion that the spatial dynamics of particular DCT peptide is also tuned by itself via inhibition (CMI) on Ca_V_1 channels or Ca^2+^ influx (Figure 8H). The clear difference between CCT_D__V/A and CCT_D_ resembled that between CCAT_C_ and CCT_C_, ending up with distinct positions along the line that also indicate their differential effects on neurite outgrowth. Taken together, our data consolidate the critical role of overall CMI: to directly gauge the level of Ca_V_1 activities that control cytonuclear distribution of DCT peptides and thus the subsequent dendritogenesis signaling, which is subject to cytosolic Ca^2+^/Ca_V_1-dependent and nuclear DCT/TF-mediated regulations.

**Figure 8.**
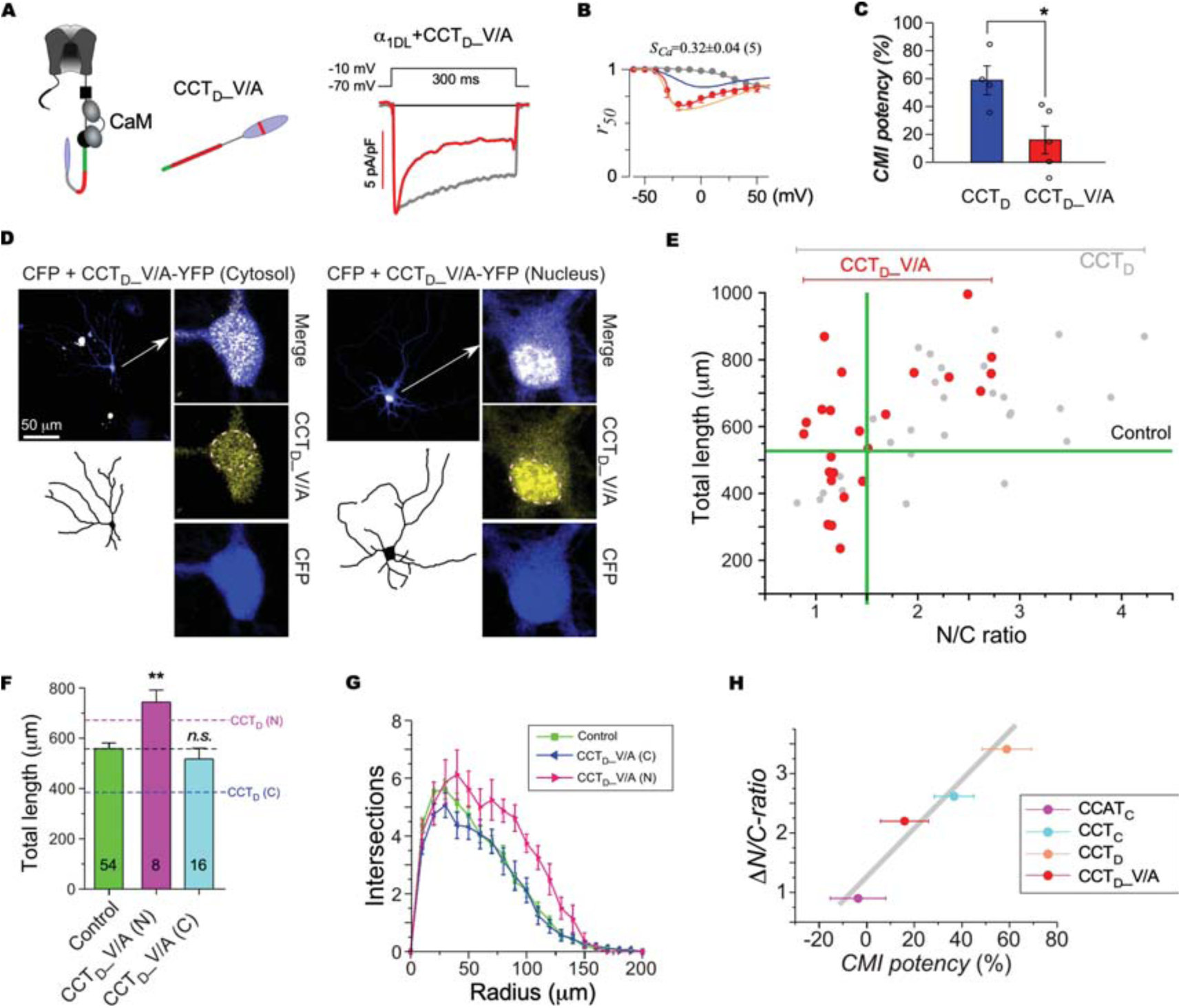
Critical roles of CMI confirmed with CCT_D__V/A mutant. (**A-C**) Cartoon illustration and representative current traces of CCT_D__V/A on α_1DL_ (**A**); and the effects of the key V/A mutation on the profiles of inactivation (**B**) and CMI potency (**C**). In comparison with CCT_D__V/A (in red), the *r_50_* profiles by the lines in blue and faint-red represent the groups of CCT_D_ and α_1DL_ control, respectively. Original data for the control and CCT_D_ were adopted from Figure 3A. (**D**) Morphology of cortical neurons with cyto-nuclear CCT_D__V/A. By similar criteria: nuclear (N/C ratio>1.5) and cytosolic (N/C ratio<1.5), CCT_D_ neurons were classified into the two major groups of CCT_D_ (N) and CCT_D_ (C) respectively, demonstrated here with the confocal images, neurite tracing, and cyto-nuclear distribution in the similar style of Figure 6A. (**E** to **G**) Cyto-nuclear distribution of CCT_D__V/A peptides in correlation with neurite outgrowth. The scatter plot of CCT_D_ was adopted from Figure 7C as the control group to compare with CCT_D__V/A (**E**), further summarized in the context of total length (**F**) or *Sholl* analyses (**G**) for nuclear and cytosolic CCT_D__V/A. The dashed lines in (**F**) represent the averages of CCT_D_ (N) and CCT_D_ (C) from Figure 6B. (**H**) Correlation between CMI potency and cyto-nuclear distribution for each of the four peptides: CCAT_C_, CCT_C_, CCT_D_ and CCT_D__V/A. Student’s *t*-tests and one-way ANOVA followed by Dunnett for post hoc tests were used for (**C**) and (**F**), respectively (*, *p*<0.05; **, *p*<0.01). Data were shown as means±SEM.

## Discussion

In this study, we propose a dynamic tuning scheme based on systematic examinations of DCT peptides derived from Ca_V_1 channels, unveiling that each peptide could switch between cytosolic inhibitor of dendritogenesis and nuclear transcription factor of neurotrophic genes. Both signaling opponents are gauged by CMI intrinsic to each peptide, either by tuning its interaction with Ca_V_1 and subsequent Ca_V_1-dependent transcriptional signals, or by tuning its Ca^2+^-dependent cytonuclear translocation, altogether constituting a delicate tunable balance in neurons.

### Tuning scheme of signaling opponents centered with CMI of DCT/Ca_V_1

Although the neurotrophic effects of DCT peptides have been demonstrated as transcription factors in the nucleus, the overall effects of these DCT peptides on neurite outgrowth appear inconclusive in cultured cortical neurons (Figure 2). Also considering the destructive effects through peptides’ CMI, we propose a *de novo* scheme to describe the homeostatic tuning of DCT/Ca_V_1-dependent neuron morphogenesis (Figure 9). We unveil that CMI inhibits neurite outgrowth, presumably by downregulating Ca_V_1-depdendent excitation-transcription coupling which includes a series of events: activation of CaMKII, phosphorylation of CREB and transcription of neurotrophic genes (e.g. Egr1, Gjb4, β-catenin) (Impey, McCorkle et al. 2004, Wheeler, Barrett et al. 2008, Wheeler, Groth et al. 2012, Ma, Groth et al. 2014). Therefore, ranging from CCAT_C_ to CCT_D_, higher CMI potency leads to less Ca^2+^ influx via Ca_V_1, eventually causing more profound retraction. On the other hand, less Ca^2+^ influx by higher CMI potency also causes more nuclear retention due to Ca^2+^-dependent nuclear export of the peptide. Nuclear DCT peptides as transcription factors directly promote expression of neurotrophic genes, e.g. Egr1, Gjb5, Fmn (Levkovitz and Baraban 2002, Gomez-Ospina, Tsuruta et al. 2006, Chasseigneaux, Dinc et al. 2011, Sahasrabudhe, Ghate et al. 2016, Duclot and Kabbaj 2017). Notably, although cytosolic CMI and nuclear transcription apparently are not directly related, higher CMI potency is actually able to upregulate nuclear fraction of the peptide and thus its function as transcription factor. The spatial regulations by CMI potency delicately balance between retraction and outgrowth of the neurites for each DCT variant. For example, CCAT_C_ exhibits ultraweak CMI or inhibition of Ca_V_1 signals with larger fraction of cytosolic distribution, opposed to TF signals from its relatively small fraction in the nucleus. CCT_D_ has much more potent CMI but relatively less cytosolic fraction (in opposition to nuclear peptide of large fraction), the retraction and outgrowth signals also nearly balance out. For either CCAT_C_ or CCT_D_, the overall balance of signaling opponents accounts for the rather mild effects of peptides on neurite morphology. To this end, an interesting scheme of cell signaling is emerging that one signaling molecule could autonomously tune the spatial (cyto-nuclear) dynamics of its own, as the way to dynamically tune/balance its opposing roles in cell signaling, which is expected to broadly find its manifestations.

**Figure 9.**
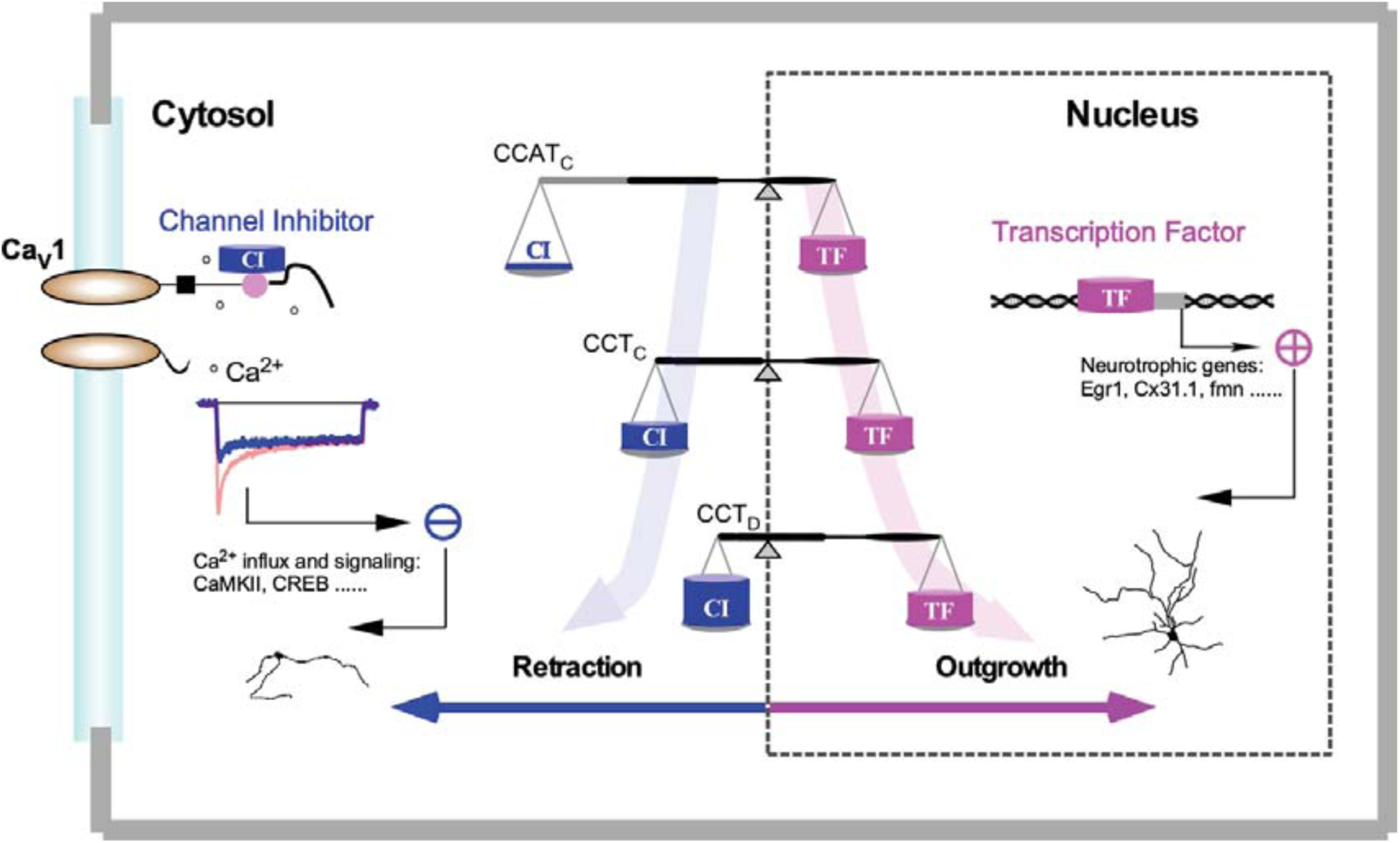
Gene regulation scheme with Ca_V_1 channels and Ca_V_1-encoded DCT peptides. DCT peptide variants act as cytosolic channel inhibitor (CI, blue) or nuclear transcription factor (TF, pink), which mediate two signaling pathways opposed to each other. DCT peptides inhibit Ca_V_1 gating and Ca^2+^ influx (CMI), thus downregulating neurotrophic genes medicated by excitation-transcription coupling. On the other hand, Ca_V_1 (through CMI) regulates cytonuclear translocation of DCT peptides, which tunes nuclear gene transcription critical to neurite outgrowth. The actual setpoint of the homeostatic balance between the above two signaling opponents is intrinsically determined by DCT peptide itself: the strength of CMI. A *de novo* scheme of Ca_V_1/DCT/CMI-centric gene regulation is emerging: two routes of transcription signals are initiated from either membrane Ca_V_1 channels or nuclear DCT peptides, subject to autonomous feedback tuning by the strength of Ca_V_1/DCT interactions (CMI).

This tuning scheme could expand onto other types of neurons. Ca_V_1 channels have been reported to express in hippocampus neurons and participate in hippocampal neurogenesis (Marschallinger, Sah et al. 2015). In fact, by overexpressing DCT peptides in cultured hippocampal neurons, the results turned out to resemble those we obtained from cortical neurons (Figure 9-figure supplement 1). It is noticeable that the average N/C ratio of CCT_D_ peptides are smaller in hippocampal neurons than in cortical neurons. Such discrepancy may be attributed to the difference between these two kinds of neurons such as their different activities (Penn, Segal et al. 2016). Overexpression of CCAT_C_ in cerebellar granule cells directly promoted neurite outgrowth even without segregating the neurons into different groups (Gomez-Ospina, Tsuruta et al. 2006), potentially also due to difference in the patterns and levels of activities in different types of neurons. Even though, we postulate that the signaling scheme we propose in this work is also applicable to cerebellar granule cells. In fact, Ca_V_1 channels are closely involved in neurite growth of cerebellar granule cells (Zhao, Lu et al. 2018) and the neurite retraction caused by CCAT_C_ΔTA (truncation of transcriptional-activation domain or PCRD_C_) (Gomez-Ospina, Tsuruta et al. 2006) is consistent with the inhibitory effects of DCT peptides (CMI) reported in this work.

### Unification of DCT/CMI tuning across Ca_V_1 family

The effects of DCTs on channel gating were considered to be drastically different among Ca_V_1 family members before this work. Here, we unify the tuning scheme of CMI by DCTs across Ca_V_1.1-1.4. In particular, DCRD_S_ and DCRD_C_ are actually able to exert substantially strong CMI, as opposed to previous observations and estimations, although indeed both are relatively less potent than DCRD_D_ and DCRD_F_. Importantly, we clarify that the discrepancy in CMI strength of different DCT is critically dependent on the difference in PCRD motifs across Ca_V_1. Future work is expected to identify the key residues on PCRD other than the arginine residues reported earlier (R1696 and R1697 of Ca_V_1.2) (Hulme, Yarov-Yarovoy et al. 2006), considering the above functional profiles and the two homologous sites of ‘RR’ for Ca_V_1.1-1.3 and ‘RQ’ in Ca_V_1.4 (Figure 2-figure supplement 1). A few residues down to the above sites, i.e., S1575/T1579 or S1700/T1704 on Ca_V_1.1 and Ca_V_1.2 respectively, may hold the key (Emrick, Sadilek et al. 2010). Activities of Ca_V_1.2 could be regulated by phosphorylation of these two sites (Fuller, Emrick et al. 2010). In contrast, the two corresponding sites on Ca_V_1.3 and Ca_V_1.4 are SQ and TE, which potentially account for their CMI profiles distinct from Ca_V_1.1 and Ca_V_1.2.

The domain containing the last 55 amino acids of DCRD_C_ from Ca_V_1.2 named CT3D (Figure 2-figure supplement 1 and Figure 3-figure supplement 1) might induce CMI on Ca_V_1.2 activities in guinea-pig ventricular myocytes (Lei, Xu et al. 2018). However, our data demonstrate that truncation of CT3D (corresponding to DCRD_F_1-66 of the first 66 amino acids) merely affects CMI (Figure 3-figure supplement 1). CT3D domains less conserved across Ca_V_1 family (than its upstream) may underlie the apparent discrepancy described above, which needs future investigations.

Despite impressive progress in Ca_V_1 structures especially by taking advantages of cryo-EM, none of these structures has acquired atomic information of DCT (Wu, Yan et al. 2015, Wu, Yan et al. 2016, Zhao, Huang et al. 2019), which awaits future structural studies targeting the relevant details (Liu, Yang et al. 2010, Liu, Yang et al. 2017). We here propose a unified scheme for DCT across Ca_V_1 family, laying the foundation for the common principles and quantitative differences among DCT variants. Such tuning scheme could predict CMI of DCT peptides even if not yet being characterized. For example, the peptide of CCT_S_ is generated by cleavage of Ca_V_1.1 in skeletal muscle (Hulme, Konoki et al. 2005). The CMI potency of CCT_S_ is predicted to be relatively strong when function with Ca_V_1.3 (PCRD_D_) and rather weak when function with Ca_V_1.1 (PCRD_S_) and Ca_V_1.2 (PCRD_C_) according to CMI tuning schemes (Figure 4), suggesting that CCT_S_ would barely change Ca^2+^ influx or channel gating of Ca_V_1.1 and Ca_V_1.2, while it should effectively inhibit Ca_V_1.3. For skeletal muscle, cytosolic inhibition is minimum for CCT_S_ which should mainly act as nuclear transcriptional regulator instead of channel inhibitor since the expression level of Ca_V_1.3 there is lower than detection limit.

### Transcriptional regulations by other Ca^2+^ channels and channel encoded-peptides

Similar to Ca_V_1-encoded peptides, Ca_V_2.1 and Ca_V_3.2 could also encode peptides targeting the nucleus to regulate gene transcription by a bicistronic mechanism, potentially conserved across the super family of voltage gated Ca^2+^ channels (Kordasiewicz, Thompson et al. 2006, Du, Wang et al. 2013, Du, Wei et al. 2019). C-terminal fragments of α_1A_ also act as transcription factors to promote neuronal development (Du, Wang et al. 2013, Du, Wei et al. 2019), resembling the function of Ca_V_1 DCT peptides. Unlike the dual roles of Ca_V_1 DCT in this study, α_1A_ CT has not been found any effect on channel gating (Lubbert, Goral et al. 2017). DCT peptides of Ca_V_1 mediate multiple regulatory pathways besides inhibition of Ca^2+^/Ca_V_1-dependent transcription via CMI in this study, e.g, by directly down-regulating Ca_V_1.2 transcription when located in the nucleus of cardiac myocytes (Schroder, Byse et al. 2009, Satin, Schroder et al. 2011). Meanwhile, reduction of Ca_V_1.2 expression might also lead to less production of DCT peptides through proteolytic cleavage. Future study is needed to cope with such complicated situations. CCAT_C_ is reported to locate in cell nucleus of cerebellum and thalamus in embryos, then exporting from nucleus to cytosol along with aging and development (Gomez-Ospina, Panagiotakos et al. 2013). We suspect that age-related changes might include perturbations of homeostatic Ca_V_1/DCT balance of signaling opponents as we propose in this study, potentially also expandable onto Ca_V_2.1 (Du, Wang et al. 2013, Du, Wei et al. 2019).

Similar to the signaling scheme that Ca_V_ channels and Ca_V_1-encoded peptides mediate concurrent but distinct pathways, TRPM7 can function as chanzyme (channel kinase) which is permeable to divalent cations while participating in receptor tyrosine kinase (RTK)-mediated pathways by its α-kinase domain (Zou, Rios et al. 2019). Communications/interactions similar to those of Ca_V_1/DCT might also exist between the channel and kinase from TRPM7. We expect that one channel gene that encodes two different signaling proteins could be manifested by more signaling complexes beyond Ca_V_1 channel/transcriptional factor, e.g., TRPM7 channel/kinase.

### Autonomous regulation of nuclear distribution of transcription factors

Ca_V_1-encoded polypeptides exhibit Ca^2+^-dependent nuclear export, potentially with the aid of its nucleus retention domain NRD (Gomez-Ospina, Tsuruta et al. 2006). The exact mechanisms that control cytonuclear transport of peptides are still unclear, which may be due to direct modifications of NRD domain or indirect modulations of the proteins that control nuclear import/export. Multiple lines of evidence including our data (Figure 7 and Figure 7-figure supplement 2) demonstrate that such Ca^2+^-dependent cytonuclear translocation is highly specific to Ca^2+^ signals via Ca_V_1 (Gomez-Ospina, Tsuruta et al. 2006). The autonomous regulation of DCT peptide through modulating Ca^2+^ by binding with its precursor protein Ca_V_1 channel may serve as a prototype potentially shared by some other transcriptional regulators. A widely-expressed transcription factor NFAT also exhibits Ca^2+^-dependent nuclear import, by exposing its NLS (nuclear localization sequence) once dephosphorylated by calcineurin (Macian 2005). A similar fashion is applicable to NFAT signaling although NFAT does not belong to any part of Ca_V_1 gene and a signaling complex with more players is involved (Oliveria, Dell’Acqua et al. 2007, Murphy, Sanderson et al. 2014, Murphy, Crosby et al. 2019, Wild, Sinnen et al. 2019). Briefly, NFAT’s nuclear localization is dynamically controlled by Ca^2+^/CaM-activated calcineurin (CN) and cAMP-dependent protein kinase A (PKA), all of which could be brought together by the adaptor AKAP75/150 onto Ca_V_1 channel. Meanwhile, channel gating is reportedly modulated by the above signaling complex. Therefore, NFAT is likely to control the spatial distribution of itself by synergistic tuning of Ca^2+^/Ca_V_1. NFAT in the nucleus could also regulate transcription of genes critical to neuronal development including morphogenesis, as additional evidence of the similarity between DCT and NFAT. For nuclear transcription factors without cytonuclear translocation, signaling proteins such as CaM or CaMKII may import into the nucleus upon Ca^2+^ (to activate CREB) (Ma, Groth et al. 2014). Meanwhile, CaM and CaMKII are evidenced to closely associate with Ca_V_1 to modulate its Ca^2+^ influx (Hudmon, Schulman et al. 2005, Wheeler, Groth et al. 2012), which invite key questions, e.g., how exactly the potential signaling complex of CaMKII/CaM and Ca_V_1 is formed, and what would be the difference among CaMKII variants in channel modulation and cytonuclear translocation, all awaiting future explorations.

### Pathophysiological implications of neurodegenerative diseases

Ca_V_1 channels are involved in a variety of neuropsychiatric and neurodegenerative diseases, such as bipolar disorder, schizophrenia, Parkinson’s disease and Alzheimer’s disease (Anekonda, Quinn et al. 2011, Hurley, Brandon et al. 2013, Smoller, Craddock et al. 2013). Ca^2+^ dysregulation through Ca_V_ channels has gained increasing support for its tight correlations with neurodegenerative diseases, known as ‘Ca^2+^ hypothesis’ (Chan, Guzman et al. 2007, Chakroborty and Stutzmann 2011). However, DCT peptides in these diseases are largely ignored despite that the amount and cytonuclear distribution of DCT peptides are actually changing with age (Gomez-Ospina, Tsuruta et al. 2006). In this regard, there are still some unresolved questions pertaining to Ca_V_1 channels and DCT peptides for a better understanding toward the pathological linkage between Ca_V_1/DCT and neurodegeneration. Exemplars of such questions include: whether and why Ca_V_1 (compared with other Ca_V_) play uniquely important roles in Ca^2+^ dysregulation and subsequent neurodegeneration; what would be the physiological significance of DCT peptides by CMI (compared with other Ca_V_1 modulators) to induce channel inhibition, and (compared with other transcription factors) to regulate nuclear gene transcription; whether DCT is prone to disease-associated mutations and how malfunctional DCT would affect healthy neurons; and eventually whether and how these dysregulations could be rescued. Notably, the expression levels of CaM are downregulated in Parkinson’s disease and Alzheimer’s disease (McLachlan, Wong et al. 1987, Hurley, Brandon et al. 2013), therefore overall inhibitory effects of DCT peptides on Ca_V_1 gating and signaling could be even more intense thus perturbing the hemostasis we propose in this study (Figure 9).

## Materials and Methods

### Molecular biology

The channel plasmids were constructed from α_1S_ (mouse Ca_V_1.1, XM_983862.1, Genbank^TM^ accession number), α_1C_ (human Ca_V_1.2 long variant, NM_199460.3), α_1DL_ (Ca_V_1.3 long variant, from human NM000720 or rat NM_017298.1), and α_1F_ (human Ca_V_1.4 NP005174). α_1DΔ_ was generated by truncation of α_1DL__V/A with a unique XbaI site following the IQ domain. For chimeric α_1DΔ_-PCRD_D_ and α_1DΔ_-DCT_F_, desired segments were PCR-amplified with SpeI and XbaI sites and cloned into aforementioned α_1DΔ_. Rat Ca_V_1.3 DHP^−^ was generated by single point mutation T1033Y (Zhang, Maximov et al. 2005) on α_1DΔ_, α_1DL_, α_1DL__V/A or α_1DΔ_-DCT_F_, respectively.

Constructs of CFP/YFP-DCRD_F_ in pcDNA3 were made with a similar process described previously (Erickson, Liang et al. 2003). Other CFP/YFP-tagged constructs were generated by replacing DCRD_F_ with appropriate PCR amplified segments, via unique NotI and XbaI sites, including YFP-DCRD_F_ truncations, CFP-DCRD_S/C/D_, YFP-preIQ_3_-IQ_D_-PCRD_S/C/D/F_, CFP/YFP-PCRD_S/C/D/F_-DCRD_F_, YFP-PCRD_C_-DCRD_C_ and YFP-CCAT_C_. DCRD_S/C/D_-YFP, CCT_C_-YFP and CCT_D_-YFP were based on another vector template of interest-YFP in pcDNA3 with the cloning sites of KpnI and NotI on 5’. Single point mutations such as CFP-DCRD_F__V/A and CCT_D__V/A-YFP were made by overlap PCR. To target DCT peptides to nucleus or cytosol, nuclear localization signal (NLS) (PKKKRKV) or nuclear export signal (NES) (LALKLAGLDIGS) was fused to N-terminus of YFP-DCT peptides by overlap PCR, to achieve NLS-YFP-CCAT_C_, NES-YFP-CCAT_C_, NLS-YFP-CCT_C_ and NES-YFP-CCT_C_.

### Dissection and culturing of cortical neurons and hippocampal neurons

Cortical neurons and hippocampal neurons were dissected from newborn ICR mice. Isolated cortex tissues or hippocampal tissues were digested with 0.25 % trypsin for 15 min at 37 °C, followed by terminating the enzymatic reaction by DMEM supplemented with 10 % FBS. The suspension of cells was sieved through a filter then centrifuged at 1000 rpm for 5 minutes. The cell pellet was resuspended in DMEM supplemented with 10% FBS and were plated on poly-D-lysine-coated 35-mm No. 0 confocal dishes (In Vitro Scientific). After 4 hours, neurons were maintained in Neurobasal medium supplemented with 2 % B27, 1 % glutaMAX-I (growth medium) for 5 days. Temperature of 37 °C with 5 % CO_2_ were controlled in the incubator. All animals were obtained from the laboratory animal research center, Tsinghua University.

Procedures involving animals have been approved by local institutional ethical committees of Tsinghua University and Beihang University.

### Transfection of cDNA constructs in HEK293 cells and cultured neurons

For electrophysiological recording, HEK293 cells (ATCC), checked by PCR with primers 5’-GGCGAATGGGTGAGTAACACG −3’ and 5’-CGGATAACGCTTGCGACCTATG −3’ to ensure free of mycoplasma contamination, were cultured in 60-mm dishes, and recombinant channels were transiently transfected according to established calcium phosphate protocol (Liu, Yang et al. 2010, Liu, Yang et al. 2017). 5 µg of cDNA encoding channel α_1_ subunit, along with 4 µg of rat brain β_2a_ (M80545) and 4 µg of rat brain α_2_δ (NM012919.2) subunits were applied to HEK293 cells. To enhance expression, cDNA for simian virus 40 T antigen (1 µg) was also co-transfected. For each additional construct, 2 µg cDNA was added. All of the above cDNA constructs were driven by a cytomegalovirus (CMV) promoter. Cells were washed with PBS 6–8 h after transfection and maintained in supplemented DMEM, then incubated for at least 48 h in a water-saturated 5% CO_2_ incubator at 37 °C before usage.

For transfection in neurons, 2 µg of cDNA encoding the desired peptides were transiently transfected by Lipofectamine 2000 (Invitrogen) for each confocal dish with a typical protocol according to the manual. Mixture of plasmids and Lipofectamine 2000 in opti-MEM was added to the Neurobasal medium for transfection. After 2 hours, neurons were maintained in Neurobasal medium supplemented with 2 % B27, 1 % glutaMAX-I for 48 hours.

For 2-hybrid 3-cube FRET experiments, HEK293 cells were cultured on confocal dishes. FRET cDNA constructs of 2 µg each were transfected by Lipofectamine 2000 for 6 hours with similar protocol. Cells were used after 24 hours.

### Whole-cell electrophysiology

Whole-cell recordings of transfected HEK293 cells were performed at room temperature (25 °C) using an Axopatch 200B amplifier (Molecular Devices). Electrodes were pulled with borosilicate glass capillaries by a programmable puller (P-1000, Sutter Instrument) and heat-polished by a microforge (MF-830, Narishige), resulting in 2-5 MΩ resistances before 70% of compensation. The internal/pipette solution contained (in mM): CsMeSO_3_, 135; CsCl_2_, 5; MgCl_2_, 1; MgATP, 4; HEPES, 5; and EGTA, 5; at 290 mOsm adjusted with glucose and at pH 7.3 adjusted with CsOH. The extracellular/bath solution contained (in mM): TEA-MeSO_3_, 135; HEPES, 10; CaCl_2_ or BaCl_2_, 10; 300 mOsm, adjusted with glucose and at pH 7.3 adjusted with TEAOH, similar to the previous protocols (Liu, Yang et al. 2017). Whole-cell currents were generated from a family of step depolarizations (−70 to +50 mV from a holding potential of −70 mV and step increase of 10 mV). Current traces were recorded at 2 kHz low-pass filtering in response to voltage steps with minimum interval of 30 s. P/8 leak subtraction was used throughout. Ca^2+^ current was normalized over different cells by cell capacitance (*C_m_*, in pF), and *I_peak_* at −10mV or *I_Ca_* (in pA/pF) serves as the major index for VGA.

### 2-hybrid 3-cube FRET

2-hybrid 3-cube FRET experiments were carried out with standard protocols as described previously (Erickson, Liang et al. 2003, Liu, Yang et al. 2010, Liu, Yang et al. 2017). Briefly, experiments were performed on an inverted epi-fluorescence microscope (Ti-U, Nikon), with computer-controlled filter wheels (Sutter Instrument) to coordinate with diachronic mirrors for appropriate imaging at excitation, emission and FRET channels. The filters used in the experiments were excitation: 438/24 (FF01-438/24-25, Semrock) and 480/30 (FITC, Nikon); emission: 483/32 (FF01-483/32-25, Semrock) and 535/40 (FITC, Nikon); dichroic mirrors: 458 nm (FF458-Di02-25×36, Semrock) and 505 nm (FITC, Nikon). Fluorescence images were acquired by Neo sCMOS camera (Andor Technology) and analyzed with 3^3^-FRET algorithms coded in Matlab (Mathworks), mainly based on the following formula:

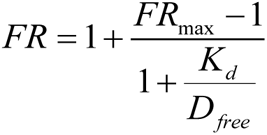

*FR_max_* represents the maximum FRET ratio, and *D_free_* denotes the equivalent free donor (CFP-tagged) concentration. *K_d_* (effective dissociation equilibrium constant) is to evaluate the binding affinity for each pair of binding partners. FRET imaging experiments were performed with HEK293 cells in Tyrode’s buffer containing 2 mM Ca^2+^.

### Confocal fluorescence imaging and analysis

Cultured neurons were transfected with CFP (to label the soma area and neurites) and DCT peptides tagged with YFP on 5^th^ day (DIV-5) and used on 7^th^ day. Neurons were loaded with Hoechst 33342 for 5 min to label the nuclei and then imaged by Zeiss LSM710 confocal Scanning Microscope. Fluorescent intensity was quantified and analyzed with ImageJ (NIH).

Calculation of nuclear intensity for YFP-tagged DCT peptides was based on the nuclear contour indicated by Hoechst 33342. Cytosolic intensity for DCT peptides was calculated by intermediate region between nucleus and plasma membrane. N/C ratio of DCT peptides was calculated by the ratio of fluorescence intensity (nuclear/cytosolic). Measurements of the total length and *Sholl* analysis for neurites were performed with Imaris 7.7.2 (Bitplane) through CFP channel. Only non-overlapping neurons were selected for analysis. Neurite tracings were depicted with Imaris 7.7.2 and further processed with Photoshop 7.0 (Adobe).

To observe the cyto-nuclear translocation of DCT peptides, neurons were pre-incubated in 5 mM [K^+^]_o_ solution (130 mM NaCl, 5 mM KCl, 1 mM MgCl_2_, 15 mM HEPES, 2 mM CaCl_2_, at 300 mOsm adjusted with glucose) and perfused with 40 mM [K^+^]_o_ solution (95 mM NaCl, 40 mM KCl, 1 mM MgCl_2_, 15 mM HEPES, 2 mM CaCl_2_, at 300 mOsm adjusted with glucose) or 5 mM [K^+^]_o_ with 50 μM Nifedipine for 0.5-1 hour, then washed out by 5 mM [K^+^]_o_ when needed. For the experiments with DHP-insensitive variants of Ca_V_1, neurons were incubated with 50 μM Nifedipine for at least 1 hour, and neurons without clear damages were selected to calculate N/C ratio for the peptides.

Analyses on neurite morphology and cyto-nuclear translocation were performed over cultured neurons of two or three different batches, adding up to the total number for each data group (20 cells or more).

### Data analysis and statistics

Data were analyzed in Matlab, OriginPro and GraphPad Prism software. Data were shown as means±SEM (Standard Error of the Mean). Unpaired Student’s *t*-test (two-tailed with criteria of significance) were performed to compare two groups. One-way ANOVA followed by Dunnett or Bonferroni for post hoc tests were performed to compare more than two groups with or without a restrictive control group, respectively. Significance *, *p*<0.05; **, *p*<0.01; ***, *p*<0.001 and *n.s.* denotes ‘not significant’.

## Acknowledgements

We thank all X-Lab members for discussions and help. This work is supported by grants from Natural Science Foundation of China (NSFC 21778034 and 81971728 for XDL, 11902021 for YXY) and of Beijing (BSFC 7191006 for XDL, 5204037 for YXY), China Postdoctoral Science Foundation (BX20180027 and 2018M641146 for YXY), and open fund from Laboratory for Biomedical Engineering of Ministry of Education, Zhejiang University.

## Author Contributions

XDL conceived the project; XDL, YBF, XML, SS and PW provided general help; YXY participated into all phases of the project; NL made unique contributions to the project including pilot experiments of peptide effects; YXY, ML and NL performed the experiments; YXY and XDL analyzed the data; WXL, ZY, WLH, PL, HJ and HYG provided technical assistance; and XDL and YXY wrote the paper.

## Supplementary Figures

**Figure 2-figure supplement 1.**
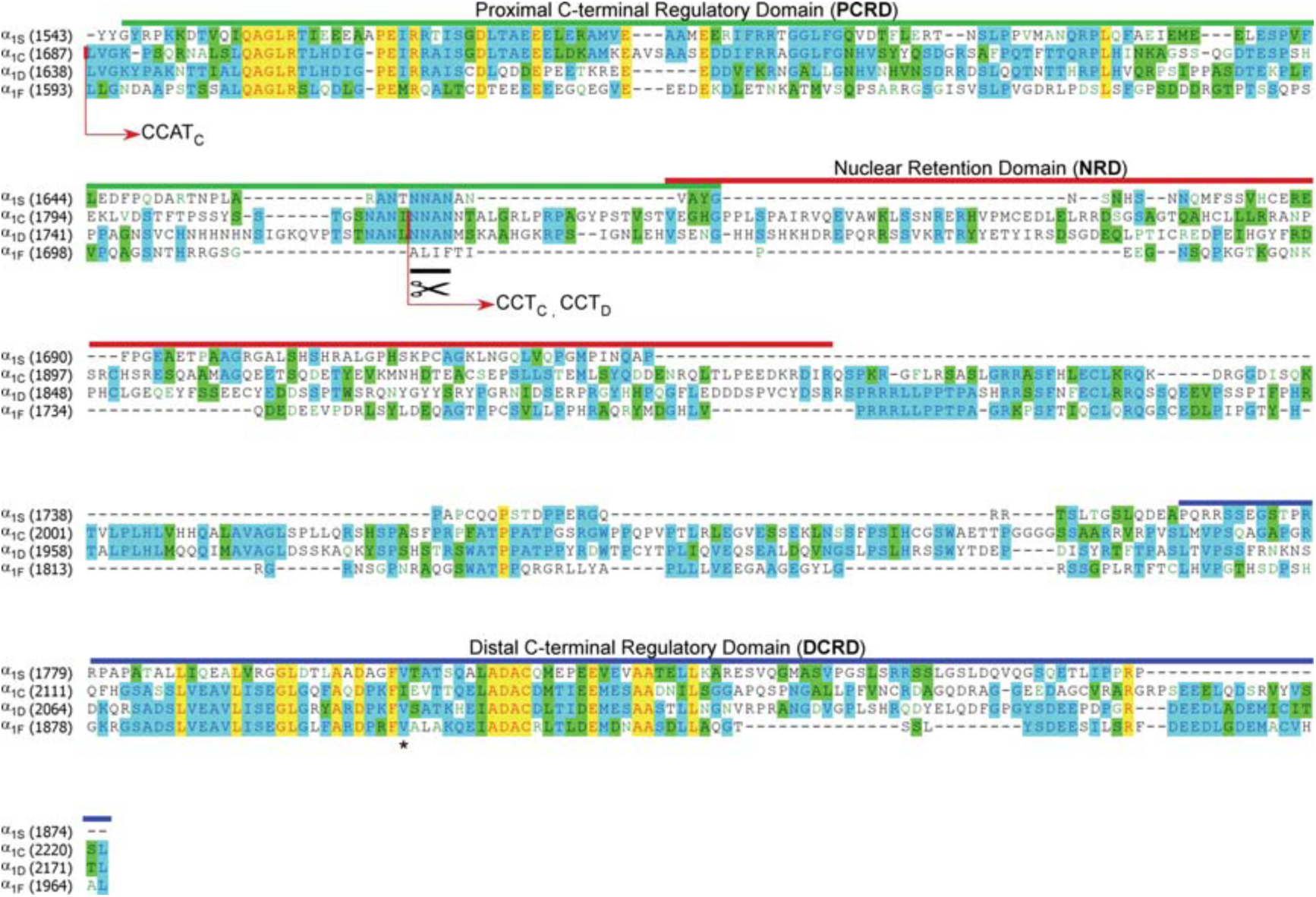
Alignment of DCT across Ca_V_1 family. The sequences of DCT were aligned for Ca_V_1.1 (α_1S_, XM_983862.1), Ca_V_1.2 (α_1C_, NM_199460.3), Ca_V_1.3 (α_1D_, NM_000720) and Ca_V_1.4 (α_1F_, NP005174), with GenBank accession numbers in parentheses. Key domains of PCRD, NRD or DCRD are indicated with solid lines of different colors; and the endogenous peptides of CCAT_C_, CCT_C_ and CCT_D_ (Gomez-Ospina, Tsuruta et al. 2006, Schroder, Byse et al. 2009, Lu, Sirish et al. 2015) are encoded by the sequences starting from the arrows (red) to the very end. Notably, PCRD is approximately overlapped with CCAT’s N-terminal transcription activation domain; and DCRD is approximately overlapped with CCAT’s C-terminal transcription activation domain (Gomez-Ospina, Tsuruta et al. 2006). Sequence homology from high to low is categorized as yellow (fully identical),cyan (partially conserved), or green (similar) and – (gapped or different). Black scissor represents the cleavage sites for potential proteolysis in the cell (Hulme, Konoki et al. 2005). Black asterisk highlights the key valine residue (i.e., unveiled by Valine to Alanine or V/A mutation) critical to DCRD functions (Liu, Yang et al. 2010).

**Figure 3-figure supplement 1.**
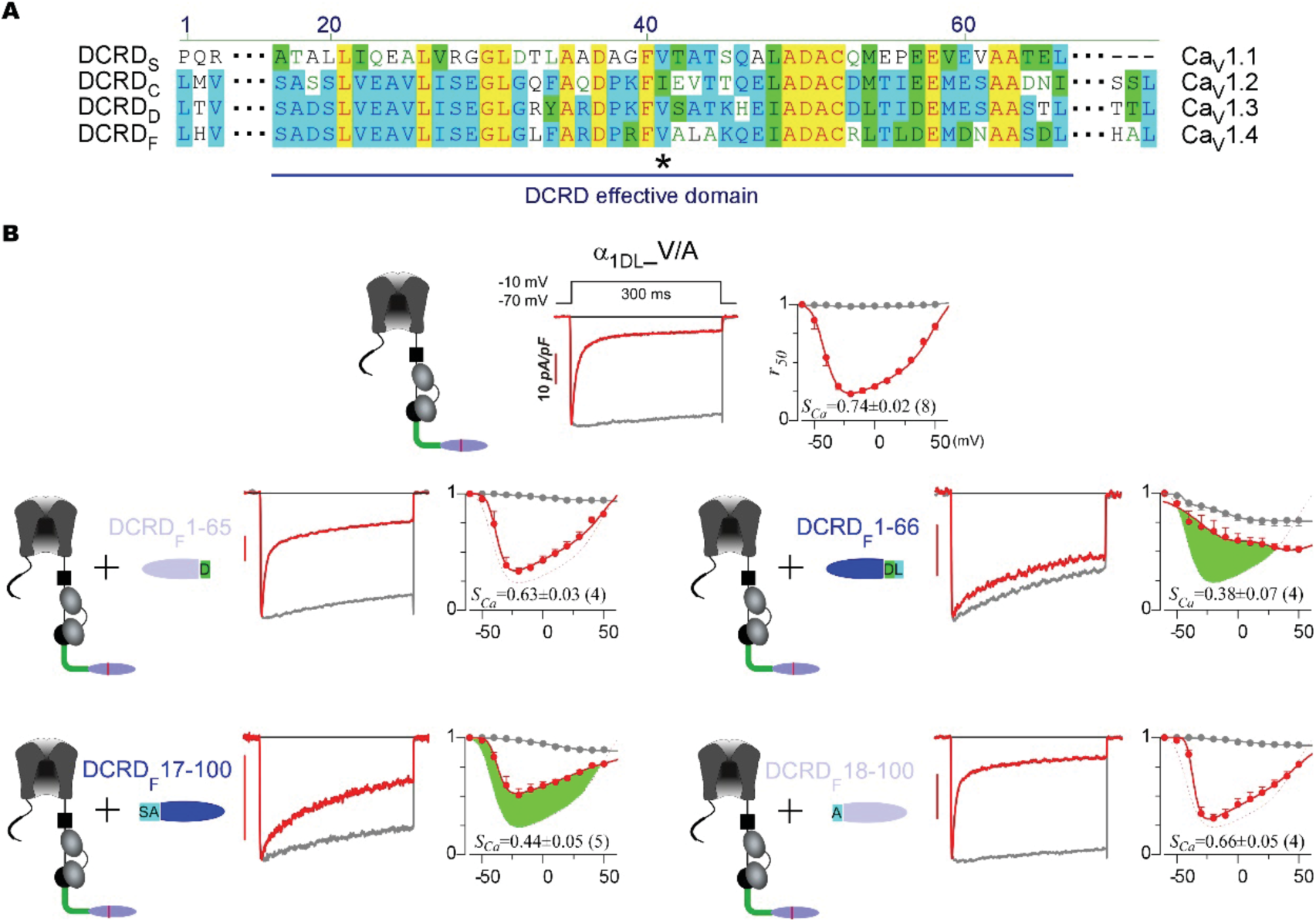
Effective motifs within DCRD across Ca_V_1 family. (**A**) Alignment of homologous DCRD subdomains across Ca_V_1 family. (**B**) α_1DL__V/A, which contains a loss-of-function mutation in DCRD domain (Liu, Yang et al. 2010), exhibited ultrastrong inactivation (strong CDI or weak CMI). In the context of α_1DL__V/A channels and DCRD_F_ peptides, the screening was conducted for the core DCRD motif. Fragments of DCRD_F_1-66 and DCRD_F_17-100 strongly attenuated CDI of α_1DL__V/A (by way of peptide CMI), whereas fragments of DCRD_F_1-65 and DCRD_F_18-100 exhibited rather weak CMI, highlighting the effective domain as DCRD_F_17-66, potentially applicable to other DCRD family members. Data were shown as means±SEM.

**Figure 4-figure supplement 1.**
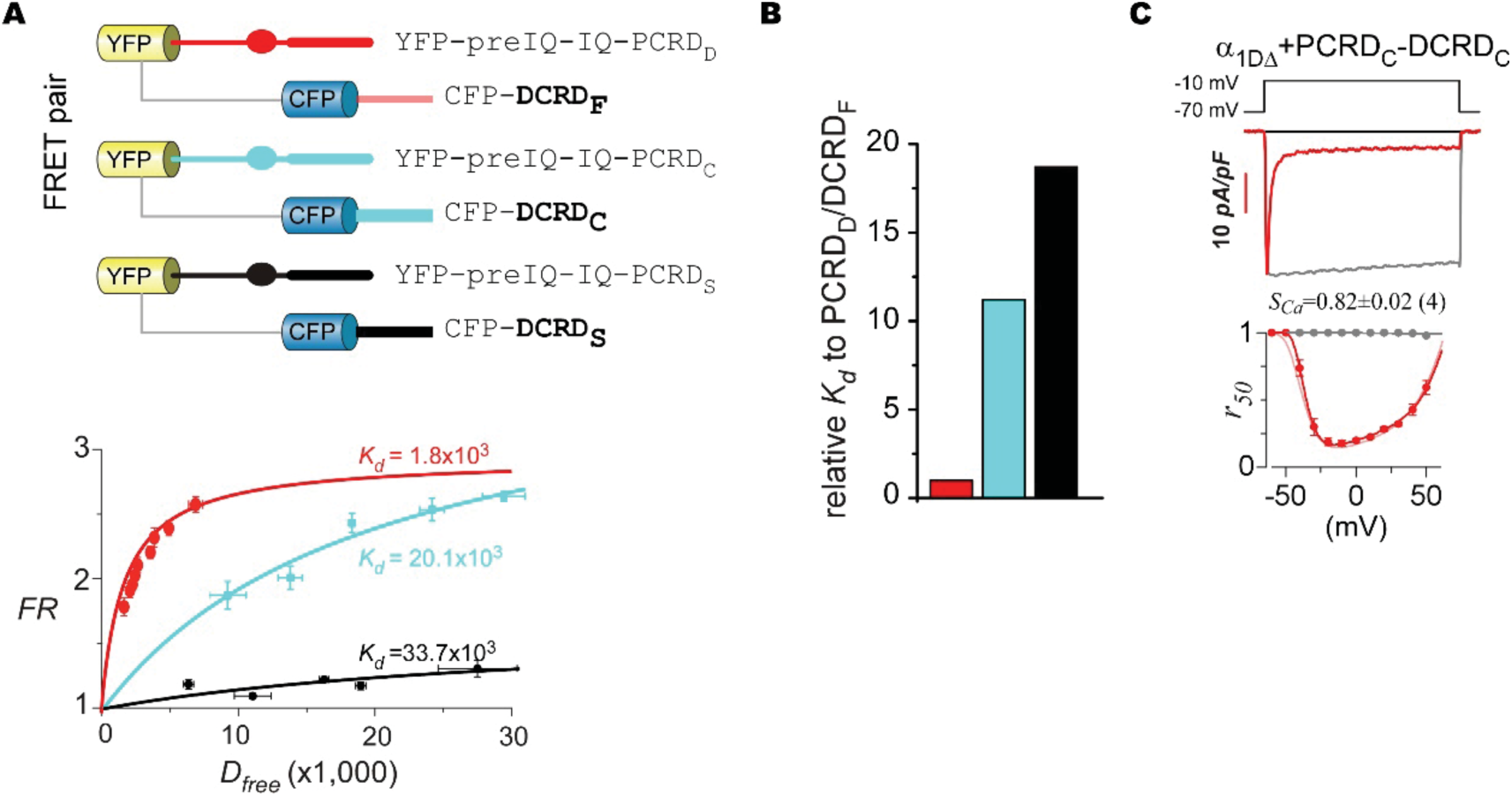
Additional binding and functional analyses for PCRD and DCRD variants from Ca_V_1.1 and Ca_V_1.2. (**A** and **B**) Binding curves for CFP-DCRD_C_ to YFP-preIQ_3_-IQ_D_-PCRD_C_ (cyan) and CFP-DCRD_S_ to YFP-preIQ_3_-IQ_D_-PCRD_S_ (black) by 2-hydrid 3-cube FRET (**A**). FRET binding between CFP-DCRD_F_ and YFP-preIQ_3_-IQ_D_-PCRD_D_ (P_D_/D_F_) was concurrently performed as the reference (red) to calculate relative *K_d_*, as summarized in (**B**) for P_C_/D_C_ (cyan), P_S_/D_S_ (black) and the reference P_D_/D_F_. (**C**) Representative Ca^2+^ traces and inactivation (*r_50_*) profiles for α_1DΔ_ with PCRD_C_-DCRD_C_ (solid red), in comparison with α_1DΔ_ control (light red). Data were shown as means±SEM.

**Figure 4-figure supplement 2.**
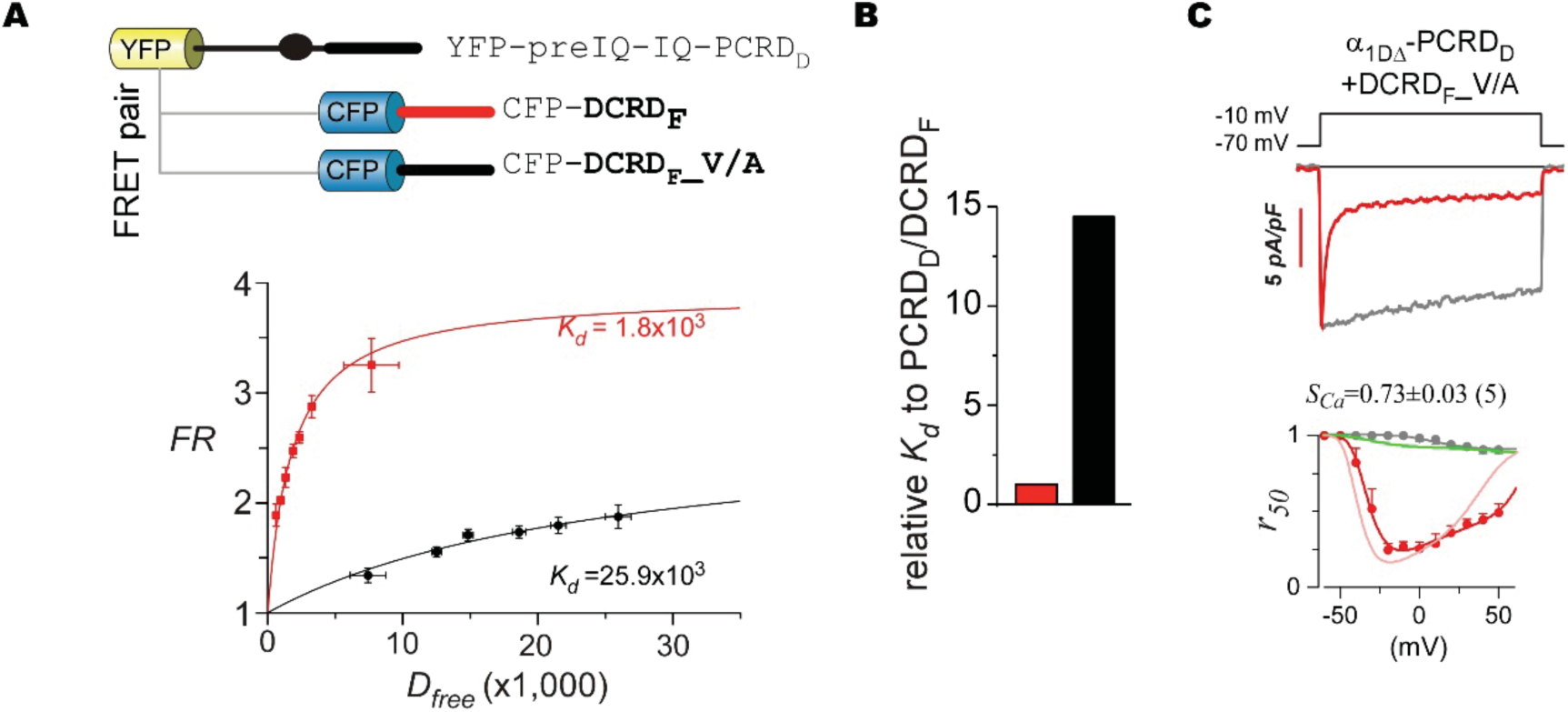
Critical V/A mutation attenuates both binding affinity and CMI potency of DCRD_F_ peptide. (**A**) Binding curve between CFP-DCRD_F__V/A (mutant DCT_F_ peptide with single-point mutation V41A) and YFP-preIQ_3_-IQ_D_-PCRD_D_ (black curve) by 2-hydrid 3-cube FRET. The red binding curve between CFP-DCRD_F_ and YFP-preIQ_3_-IQ_D_-PCRD_D_ was set as the reference. (**B**) Statistic summary of relative *K_d_* for DCRD_F__V/A to PCRD_D_ (black bar). (**C**) Representative traces and *r_50_* profiles for α_1DΔ_-PCRD_D_ co-expressed with DCRD_F__V/A. Light red curve and green curve in *r_50_* profiles represent α_1DΔ_-PCRD_D_ alone and with DCRD_F_, respectively. Data were shown as means±SEM.

**Figure 5-figure supplement 1.**
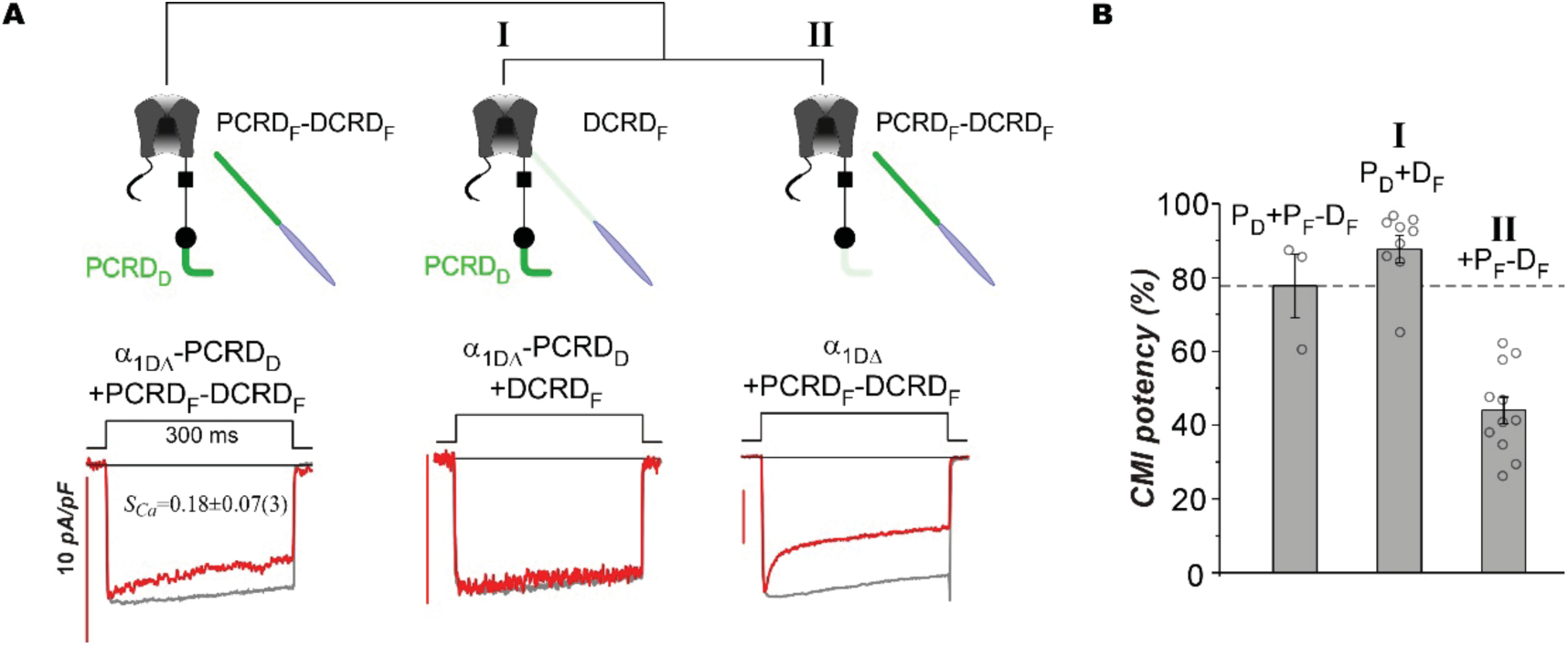
Compound CMI effects of peptide PCRD_F_-DCRD_F_ on α_1DΔ_-PCRD_D_. (**A**) Decomposition of PCRD_F_-DCRD_F_ effects on α_1DΔ_-PCRD_D_ channels. The first component (I) is the combination of peptide DCRD_F_ and channel α_1DΔ_-PCRD_D_; and the second component (II) is the combination of peptide PCRD_F_-DCRD_F_ and channel α_1DΔ_ (upper row). Representative traces corresponding to the above: P_F_-D_F_ on α_1DΔ_-PCRD_D_ and its two components (adopted from Figure 4D and Figure 4F, respectively). (**B**) Statistic summary of CMI potency. The dashed line is to compare the compound effects of α_1DΔ_-P_D_+P_F_-D_F_ with the two components: I, α_1DΔ_-P_D_+D_F_ (or P_D_+D_F_); and II, α_1DΔ_+P_F_-D_F_ (or +P_F_-D_F_). Data were shown as means±SEM.

**Figure 7-figure supplement 1.**
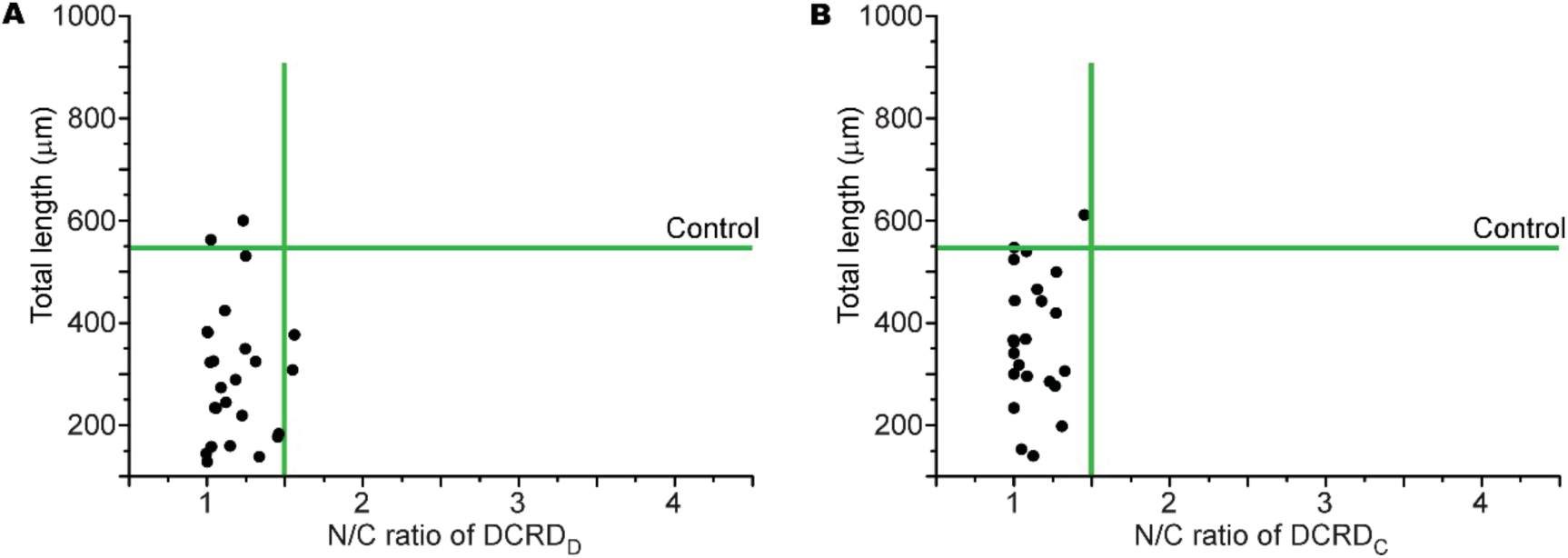
Cyto-nuclear distribution of DCRD in correlation with the total neurite length. Scatter diagrams of N/C ratio of DCRD_D_ (**A**) or DCRD_C_ (**B**) in correlation with total length of cortical neurons. Horizontal and vertical lines in green represent the control group of neurons with GFP.

**Figure 7-figure supplement 2.**
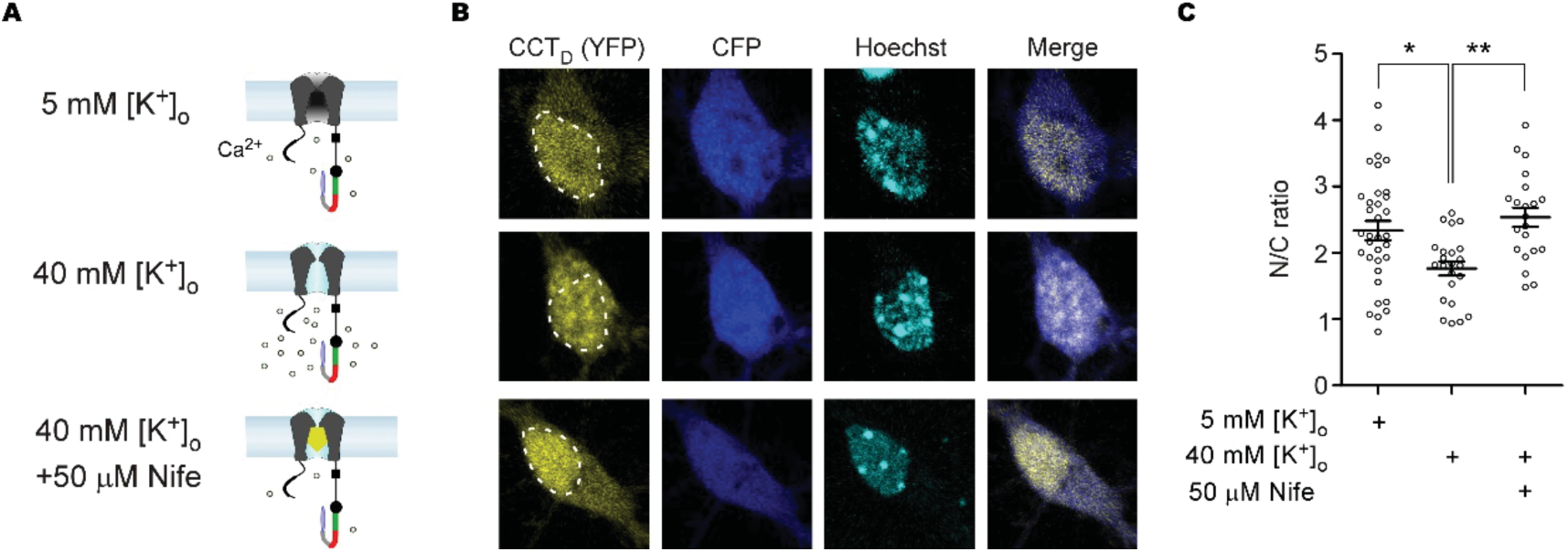
Ca_V_1 influx in response to by high potassium mediates dynamic cytonuclear translocation of DCT. (**A**) Illustration of Ca_V_1 under different conditions. Cortical neurons were maintained in 5 mM [K^+^]_o_ solutions as the basal condition (control, top), and then stimulated with 40 mM [K^+^]_o_ to induce Ca^2+^ influx (stimulated, middle). 50 μM nifedipine (Nife) blocked Ca^2+^ influx via Ca_V_1 under the conditions of 40 mM [K^+^]_o_ (blocked, bottom). (**B**) Representative confocal images of YFP (CCT_D_ distribution), CFP (soma contour), Hoechst (nuclear envelop) and YFP/CFP merge for the three groups in (**A**). (**C**) Cytonuclear translocations indexed with N/C ratio values. N/C ratio of the stimulated group was significantly suppressed when compared with the control group; whereas blockage of Ca_V_1 channels prevented the above change in N/C ratio. One-way ANOVA followed by Bonferroni for post hoc tests were used for (**C**) (*, *p*<0.05; **, *p*<0.01). Data were shown as means±SEM.

**Figure 9-figure supplement 1.**
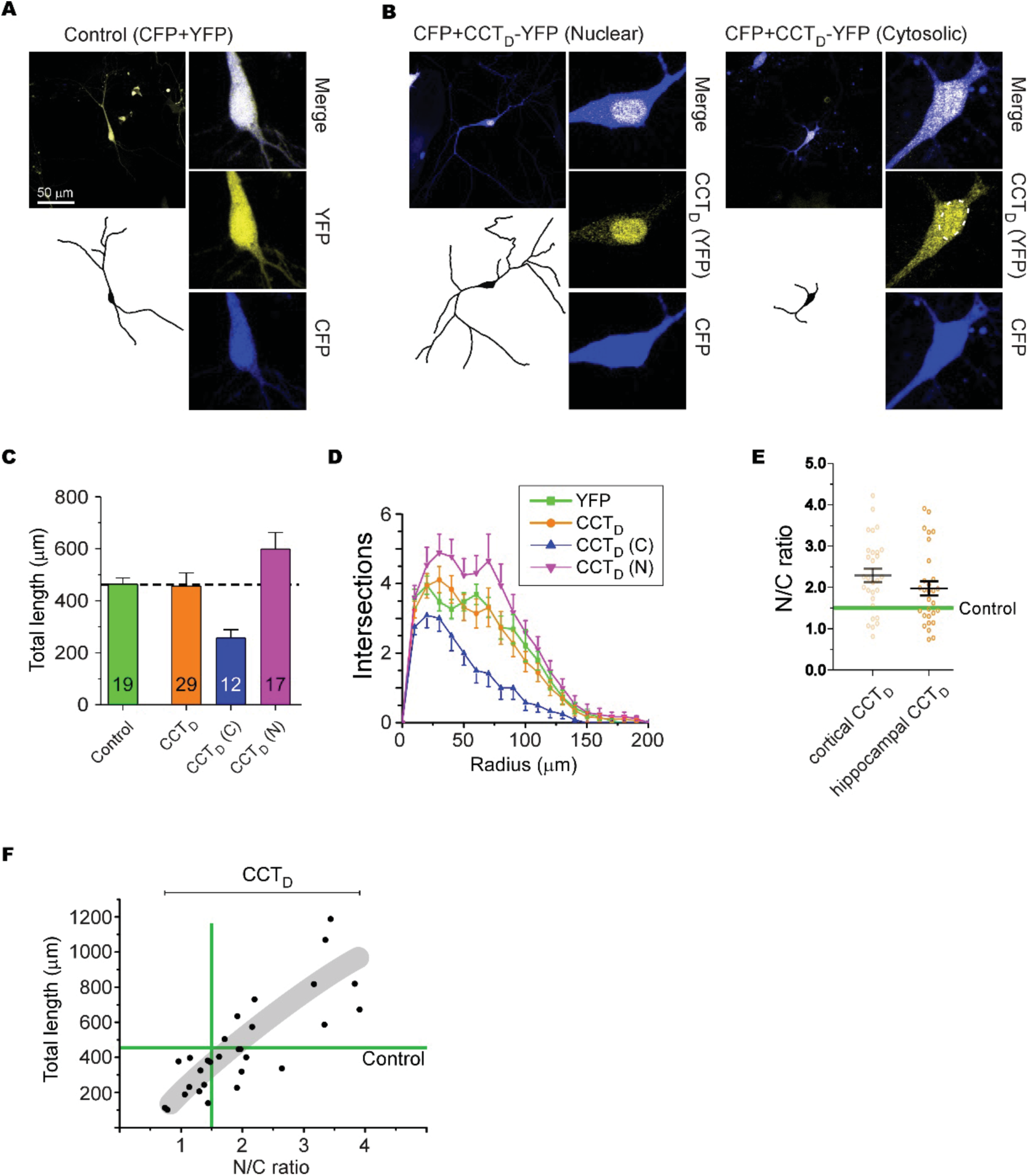
Behaviors and effects of DCT peptides in hippocampal neurons. (**A** and **B**) Cultured hippocampal neurons from newborn ICR mice (DIV-7) expressing CFP and CCT_D_-YFP were imaged with confocal microscopy. Neurons with YFP serve as control group (**A**). Neurons with CCT_D_ (**B**) were classified into the two major groups of nuclear (N) with the criteria of N/C ratio>1.5 and cytosolic (C) with the criteria of N/C ratio<1.5, respectively. Confocal images, neurite tracings, and cyto-nuclear distributions are presented in similar fashions to Figure 6A. (**C** and **D**) Total neurite length (**C**) and *Sholl* analyses (**D**) for the groups of control (YFP), total CCT_D_, CCT_D_ (C) and CCT_D_ (N). (**F**) Statistical summary of N/C ratio for cortical CCT_D_ versus hippocampal CCT_D_. Original data for cortical CCT_D_ were adopted from Figure 2E. (**F**) Cyto-nuclear distribution of CCT_D_ in correlation with neurite outgrowth. Data were shown as means±SEM.

